# Lineage-Specific Neofunctionalization of Polyacetylene-Directing UGT76 Glycosyltransferases in Campanulaceae

**DOI:** 10.64898/2026.07.02.735953

**Authors:** Jiadong Hu, Xue Cao, Min He, Chunhui Wang, Peng Di, Shanshan Chen, Chen Zhang, Ying Xiao, Renjun Mao, Wei Sun, Wansheng Chen, Shi Qiu

**Author notes:** Corresponding author: *E-mail address* (S. Qiu), (W.S. Chen), (W. Sun), (R.J. Mao). These authors contribute equally to this work (Jiadong Hu, Xue Cao, Min He, Chunhui Wang).

## Abstract

Polyacetylene glycosides exhibit notable pharmacological activities, yet the glycosyltransferases acting on their polyacetylene scaffolds remain unknown. Here we report a telomere-to-telomere genome assembly of *Codonopsis pilosula* and, guided by spatial metabolomics, characterize three UDP-glycosyltransferases: CpUGT76BG1 and CpUGT76BG2 catalyze the direct glycosylation of lobetyol to lobetyolin, while CpUGT94BY2 performs subsequent sugar-sugar coupling to produce lobetyolinin, with each activity confirmed by in planta overexpression. Structural modeling reveals that CpUGT76BG1 and CpUGT76BG2 employ a deep hydrophobic tunnel to fully encase the linear polyacetylene chain, a binding architecture distinct from the shallow pockets used by canonical plant UGTs for planar aromatic substrates. Ancestral sequence reconstruction across eleven nodes partitions the UGT76 lineage into three functionally distinct evolutionary stages, tracing the trajectory from an ancestral shallow pocket to this specialized deep architecture. These findings establish the key glycosylation steps of polyacetylene glycoside biosynthesis, define a tunnel-based paradigm for non-planar substrate recognition, and reveal how tandem duplication-driven active site remodeling generates metabolic novelty.

## Introduction

Plants are diverse producers of specialized metabolites, with lineage-specific biosynthetic pathways generating an extraordinary chemical diversity that underpins ecological adaptation and medicinal utility^1, 2^. Among these, polyacetylenes, constitute a structurally distinctive class defined by one or more carbon-carbon triple bonds, with approximately 1,400 polyacetylenoids isolated predominantly from the Asteraceae, Campanulaceae, and Apiaceae families^3–5^. Substantial pharmacological evidence supports their therapeutic potential, including antitumor, immunomodulatory, neuroprotective, and antidepressant activities^5^. Polyacetylene glycosides represent a structurally distinct subgroup, defined by carbohydrate units attached to polyacetylene alcohols via acetal linkages. This conjugation combines a hydrophobic, achiral polyacetylene core with hydrophilic, chiral sugar moieties, thereby enhancing aqueous solubility is hypothesized to facilitate intercellular and long-distance transport, while simultaneously stabilizing the reactive polyacetylene backbone to preserve bioactivity until enzymatic hydrolysis releases the active scaffold^6, 7^. A rich diversity of over 150 such polyacetylene glycosides have been identified across plant families, with notable abundance in the Campanulaceae^8^, a family comprising approximately 2400 species including medicinal herbs such as *Platycodon grandiflorus*^9^, *Codonopsis pilosula*^10^, and *Lobelia chinensis*^11^. Among these, *C. pilosula* is widely utilized in herbal medicine and as a dietary supplement, with its C-14 polyacetylene glycosides, such as lobetyolin and lobetyolinin, exhibit notable pharmacological effects including antitumor, immunomodulatory, and anti-inflammatory activities^7, 12, 13^. Despite their broad structural variety and significant bioactivities, the UDP-glycosyltransferases (UGTs) responsible for the direct glycosylation of polyacetylene scaffolds have remained entirely uncharacterized, a significant gap that limits both mechanistic understanding of polyacetylene metabolism and biotechnological access to these bioactive compounds.

UGTs play a central role in the diversification of plant specialized metabolism by modulating the solubility, stability, and bioactivity of numerous metabolites^14, 15^. The expansion and functional divergence of UGT families are primarily driven by gene duplication events, including whole-genome duplication (WGD) and tandem duplication, followed by neofunctionalization that enables the emergence of novel substrate specificities^16, 17^. Well-documented examples include the WGD- and tandem duplication-mediated functional diversification of UGTs in tea plants (*Camellia sinensis*), which contributed to flavan-3-*O*-glycosylation and likely underlies local adaptation^18^, as well as the duplication of UGTs along with serine carboxypeptidase-like (SCPL) genes that facilitated the biosynthesis of leonurine in motherwort (*Leonurus japonicus*)^19^. Despite the broad substrate range of characterized UGTs, which spans flavonoids, alkaloids, terpenoids, and phytohormones, the direct glycosylation of linear, entirely hydrophobic polyacetylene chains represents a fundamentally distinct catalytic challenge. Canonical UGT substrates such as flavonoids and phenylpropanoids are planar, aromatic molecules that engage the binding pocket through π-π stacking and directional hydrogen bonds; polyacetylenes, by contrast, are non-planar, conformationally rigid due to their carbon-carbon triple bonds, and lack polar functional groups beyond the terminal hydroxyl that serves as the glycosylation site. Accommodating such a substrate requires a binding architecture that can thread and stabilize an extended alkyne chain rather than envelop a planar aromatic ring. Elucidating these enzymes is essential not only to complete the biosynthetic pathway of polyacetylene glycosides but also to understand how UGT binding pockets can be remodeled to accommodate substrates that depart radically from the canonical structural paradigm.

Recent advances in whole-genome sequencing have greatly expanded genomic resources for medicinal plants^20, 21^, overcoming limitations associated with transcriptome-based approaches, which often obscure gene cluster architecture, collinearity, and duplication events relevant to specialized metabolism^22^. Nevertheless, genomic data alone frequently falls short in identifying key biosynthetic enzymes. Integrating spatial metabolomics with gene expression profiling provides a powerful alternative strategy. Matrix-assisted laser desorption/ionization mass spectrometry imaging enables high-sensitivity, label-free visualization of metabolite distribution within plant tissues, facilitating direct correlation between metabolic accumulation and transcriptional regulation^23^. Concurrently, evolutionary genomics offers insights into the diversification, functional divergence, and genome evolution of UGT families^24^. Together, these approaches constitute a systematic pipeline for enzyme discovery. Applied to *C. pilosula*, this multimodal strategy is uniquely suited to resolving the long-standing question of polyacetylene glycoside biosynthesis.

In this study, we employed an integrated multi-omics approach combining genomic, transcriptomic, spatial metabolomic, phylogenetic, and evolutionary analyses to elucidate the glycosylation processes underlying the biosynthesis of lobetyolin and lobetyolinin. We generated a high-quality telomere-to-telomere (T2T) genome assembly of *C. pilosula* and functionally characterized two UGTs, CpUGT76BG1 and CpUGT76BG2, that catalyze the glycosylation of lobetyol to form lobetyolin. Furthermore, we identified a di-*O*-glycosyltransferase, CpUGT94BY2, responsible for the sugar-sugar coupling step in lobetyolinin production. Molecular docking revealed that CpUGT76BG1 and CpUGT76BG2 employ a deep hydrophobic tunnel architecture to envelop the linear polyacetylene chain, a substrate-binding strategy that is structurally distinct from the open, shallow pocket utilized by the closest crystallographically characterized paralog within the UGT76 family. Ancestral sequence reconstruction traced the neofunctionalization of the UGT76 family, shedding light on the evolutionary mechanisms driving tandem duplications in Campanulaceae. These findings establish the polyacetylene glycosylation pathway at the enzymatic level, define the structural determinants of linear alkyne chain recognition by plant UGTs, and provide a molecular framework for understanding how tandem duplication-driven active site remodeling generates metabolic novelty in plant specialized metabolism.

## Results

### T2T-level genome assembly and phylogenomic analysis of the *C. pilosula* genome

We assembled a high-quality T2T genome of *C. pilosula* (designated CP2.0) to establish a reference for functional genomics. K-mer frequency analysis estimated a genome size of 1.54 Gb and a heterozygosity rate of 0.845% (Supplementary Fig. 1). Deep sequencing data were generated, including PacBio HiFi long reads (50 × coverage), Oxford Nanopore Technologies (ONT) ultra-long reads (N50 > 100 kb; 52 × coverage), and Hi-C data (120 × coverage). Using hifiasm and NextDenovo, we assembled the reads into contigs, which were subsequently scaffolded into 8 chromosomes using Hi-C interaction data processed with Juicer (Supplementary Fig. 2, Supplementary Data 1). Gaps in the chromosome-scale assembly were filled using HiFi and ONT reads along with unplaced contigs. The final CP2.0 assembly spans 949.53 Mb with a contig N50 of 120.96 Mb. Rigorous assessment through read mapping, Hi-C interaction maps, LAI, and QV metrics confirmed high assembly integrity and accuracy. Chromosome 6 was resolved to T2T completeness, and the full assembly contains only 16 gaps and 14 telomeres, achieving a BUSCO completeness of 97.4% (Fig. 1A, Supplementary Fig. 2, Supplementary Data 1–3).

**Figure 1.**
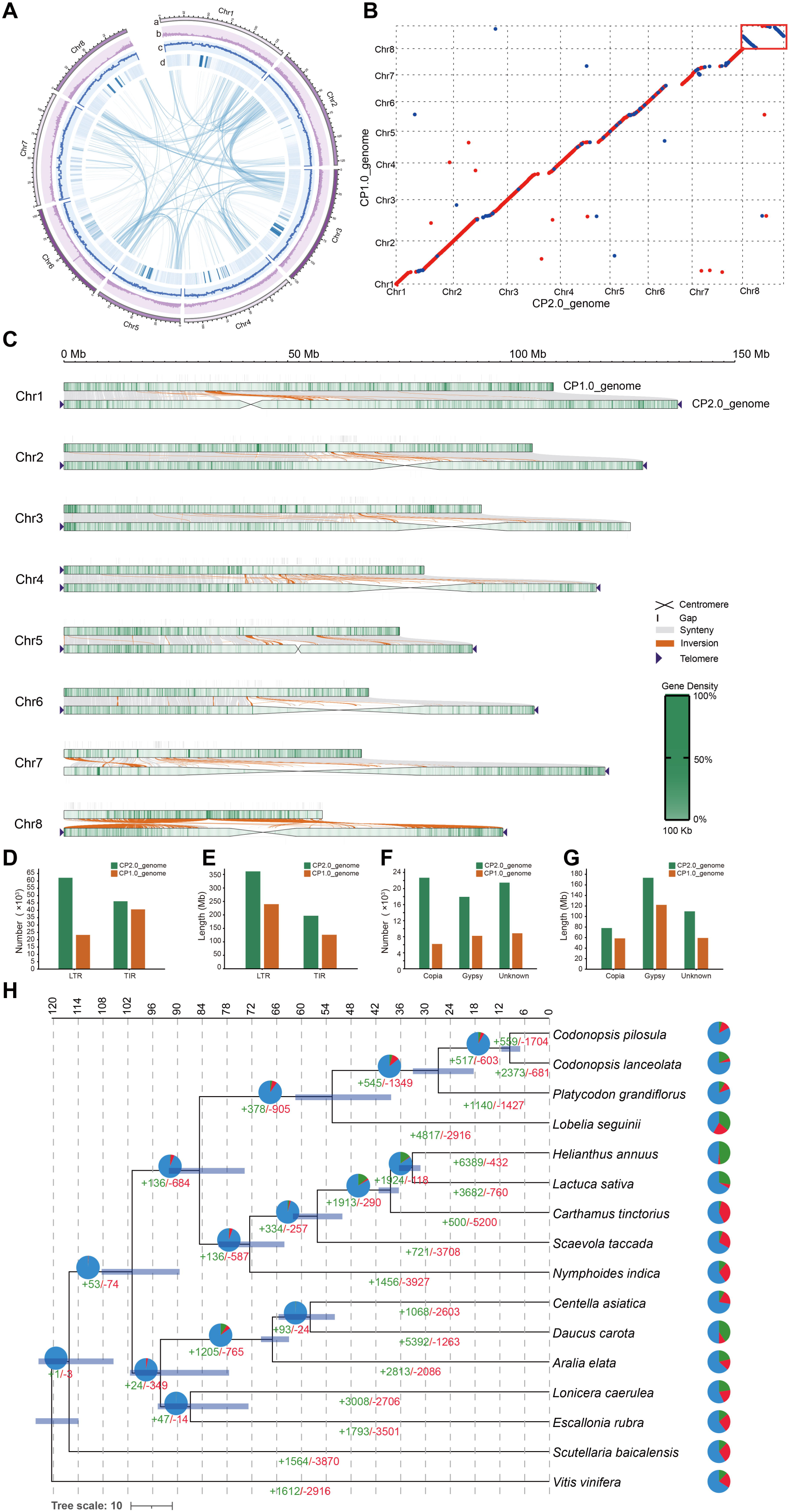
Telomere-to-telomere (T2T)-level assembly and analysis of the CP2.0 genome (*C. pilosula*). (**A**) From outer to inner rings: pseudo-chromosomes (scale in Mb), gene density, GC content, repeat density, and genome-wide syntenic blocks (colored ribbons); window size, 100 kb. (**B–C**) Collinearity of chromosomes and genome collinearity analysis between CP1.0 and CP2.0. (**D–G**) TE comparison between CP1.0 and CP2.0. (**H**) Expansion and contraction of gene families among 16 plant genomes. Scale bar, divergence time in million years ago (MYA). Transparent blue bars at nodes indicate 95% Highest Posterior Density (HPD) intervals for divergence time estimates. Branch numbers specify the exact count of gene families that have undergone expansion (green, ‘+’) or contraction (red, ‘-’). Pie charts at nodes and species tips show the proportion of gene families that have expanded (green), contracted (red), or remained stable/conserved (blue).

Gene annotation was performed using transcriptomic data from root, stem, leaf, and flower tissues, identifying 30,663 expressed genes distributed across the 8 chromosomes (Fig. 1A). The annotated gene set reached 96.3% BUSCO completeness (Supplementary Data 4-5). A total of 147 UGT genes were identified, a count consistent with other plant species (Supplementary Fig. 3). The CP2.0 assembly closed 841 of the 857 gaps present in the previous CP1.0 version^25^, adding 270.33 Mb of sequence largely located in centromeric regions, with chromosome 7 containing the largest centromere (54.19 Mb; Supplementary Data 6–8). The updated assembly corrected two large inversions on chromosome 8 and several smaller structural errors, yielding a more accurate genome architecture (Figs. 1B, C; Supplementary Fig. 2). Transposable element (TE) annotation revealed a > 15-fold increase in total TE length compared to CP1.0 (724.93 Mb vs. 46.46 Mb; Figs. 1D–1G, Supplementary Data 9), with substantial gains in LTR and TIR superfamilies. Notably, Copia-type LTRs now outnumber Gypsy-types, indicating that earlier assemblies had missed Copia-rich centromeric regions (Supplementary Data 9). These results establish CP2.0 as a superior genomic resource for Campanulaceae research.

To reconstruct the evolutionary history of Campanulaceae, we analyzed the CP2.0 genome alongside related species. *Ks* (Synonymous substitution rate) distribution analysis confirmed that *C. pilosula*, *P. grandiflorus*, *Codonopsis lanceolata*, and *Lobelia seguinii* all retain the core eudicot γ whole-genome duplication (WGD), with *L. seguinii* additionally showing a species-specific recent WGD (Supplementary Fig. 4). Phylogenomic analysis of 2,705 single-copy orthologs from 16 species identified 559 significantly expanded and 1,704 contracted gene families in *C. pilosula* (Fig. 1H). Functional enrichment analysis (KEGG and GO) indicated that expanded and contracted gene families across Campanulaceae species are significantly enriched in pathways related to fatty acid oxidation, glycosyltransferase activity, flavonoid biosynthesis, and monoterpenoid biosynthesis (Supplementary Fig. 5). The concerted expansion of these pathways provides precursors and catalytic machinery for the assembly and decoration of polyacetylene scaffolds.

### Spatial mapping of polyacetylene glycosides localization in *C. pilosula*

Leveraging this genomic foundation, we next sought to spatially resolve the biosynthesis of polyacetylene glycosides in *C. pilosula*. Using matrix-assisted laser desorption/ionization mass spectrometry imaging (MALDI-MSI) on root cross-sections in positive-ion mode, we visualized the tissue-level distributions of the polyacetylene scaffold lobetyol (aglycone) and its glycosylated derivatives, lobetyolin (monoglycoside), lobetyolinin (diglycoside), and pratianlin B (triglycoside) (Fig. 2). The glycosylated products showed strict accumulation in the periderm, whereas lobetyol was detected uniformly across all root tissues. Overlay images combining lobetyolin, lobetyolinin, and pratianlin B (Overlay-1 in Fig. 2), further emphasized the compartmentalized nature of polyacetylene glycoside biosynthesis, with end products almost exclusively restricted to the periderm (Fig. 2). This spatial patterning strongly implicates tissue-specific UGTs in the glycosylation of polyacetylene scaffolds.

**Figure 2.**
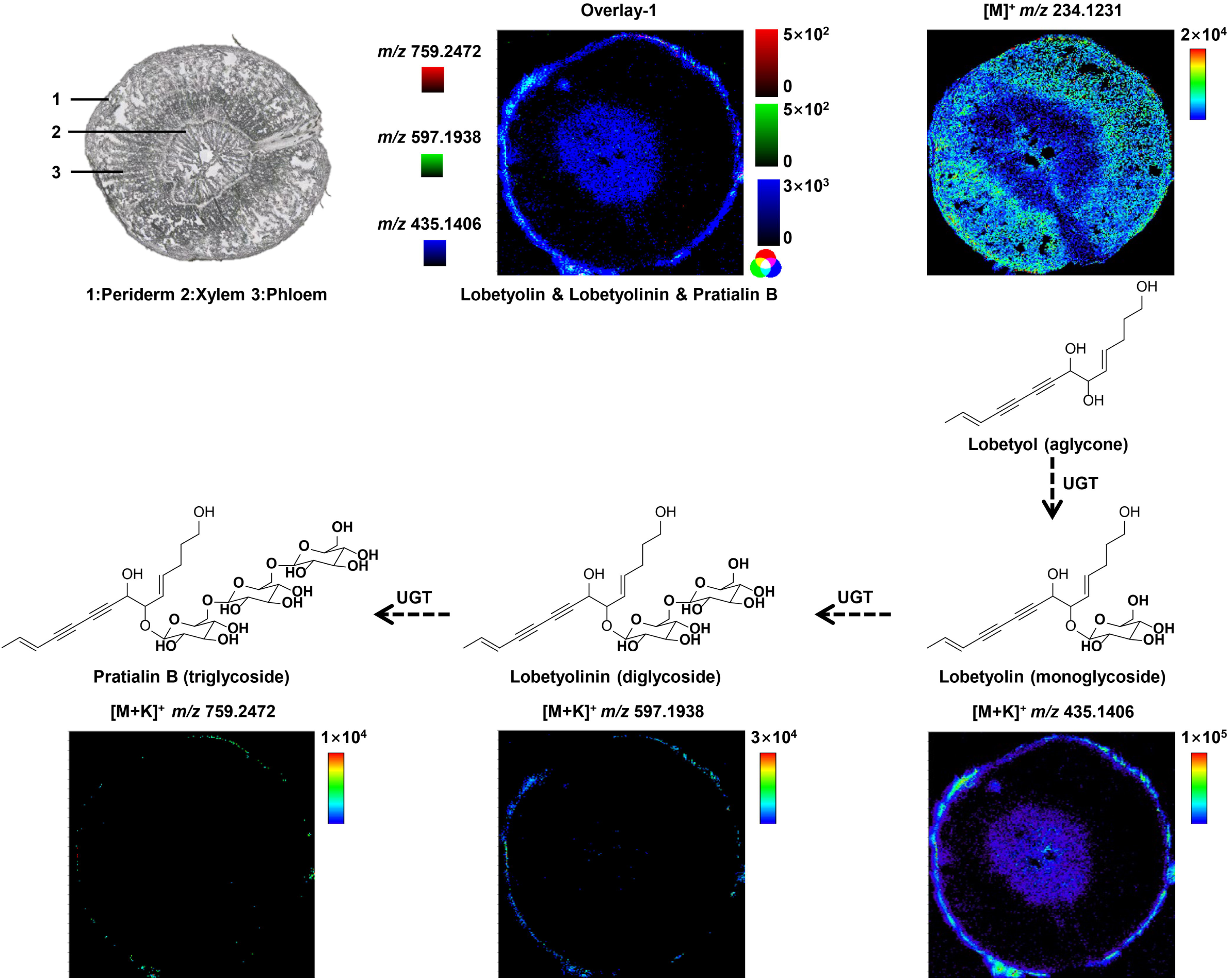
Spatial heterogeneity of polyacetylene glycosides in *C. pilosula*. An optical image of a root section is shown alongside MALDI mass spectrometry images depicting the spatial distribution of selected major ions. These include the polyacetylene scaffold lobetyol (aglycone) detected as [M]^+^at *m/z* 234.12, as well as the glycosylated derivatives lobetyolin (monoglycoside) at *m/z* 435.14 [M+K]^+^, lobetyolinin (diglycoside) at *m/z* 597.19 [M+K]^+^, and pratianlin B (triglycoside) at *m/z* 759.25 [M+K]^+^. The overlaid image (overlay-1) integrates the spatial patterns of the polyacetylene glycosides lobetyolin, lobetyolinin, and pratianlin B, further illustrating their compartmentalized biosynthesis within the root tissue. Chemical structures of lobetyol (aglycone), lobetyolin (monoglycoside), lobetyolinin (diglycoside), and pratianlin B (triglycoside) are displayed adjacent to the corresponding MALDI-MSI panels.

To quantitatively validate these spatial trends, we performed LC–MS on dissected periderm, phloem, and xylem tissues of *C. pilosula* roots. Principal component analysis of the metabolome data showed clear separation among tissue types (Supplementary Fig. 6), corroborating the imaging results. Through partial least squares-discriminant analysis, we identified lobetyolin and several alkaloids as differentially accumulated metabolites enriched in the periderm (Supplementary Figs. 7, 8, Supplementary Data 10-12). Relative quantitative analysis further confirmed that polyacetylene glycosides, including lobetyolin, lobetyolinin, and pratianlin B, were present at significantly higher levels in the periderm than in the xylem (Supplementary Fig. 8), consistent with MALDI-MSI observations. Together, these integrated findings establish the periderm as the primary biosynthetic site for polyacetylene glycosides and provided a targeted tissue-specific transcriptomic strategy, directly guided by the spatial metabolomics mapping, for identifying periderm-localized UGT candidates.

### Functional characterization of polyacetylene-specific UGTs

Guided by the spatial metabolite profiling, we conducted tissue-specific RNA sequencing of periderm, phloem, and xylem to identify UGTs potentially involved in polyacetylene glycoside biosynthesis. From a total of 147 UGT transcripts identified in the CP2.0 genome (Supplementary Fig. 3), 18 candidates showing periderm-enriched expression were selected for further analysis (Fig. 3A, Supplementary Data 20). These candidates were cloned into pET-28a vector and heterologously expressed in the *E. coli* (BL21) strain. Recombinant proteins were affinity-purified using Ni-NTA agarose for subsequent enzymatic assays (Supplementary Fig. 9).

**Figure 3.**
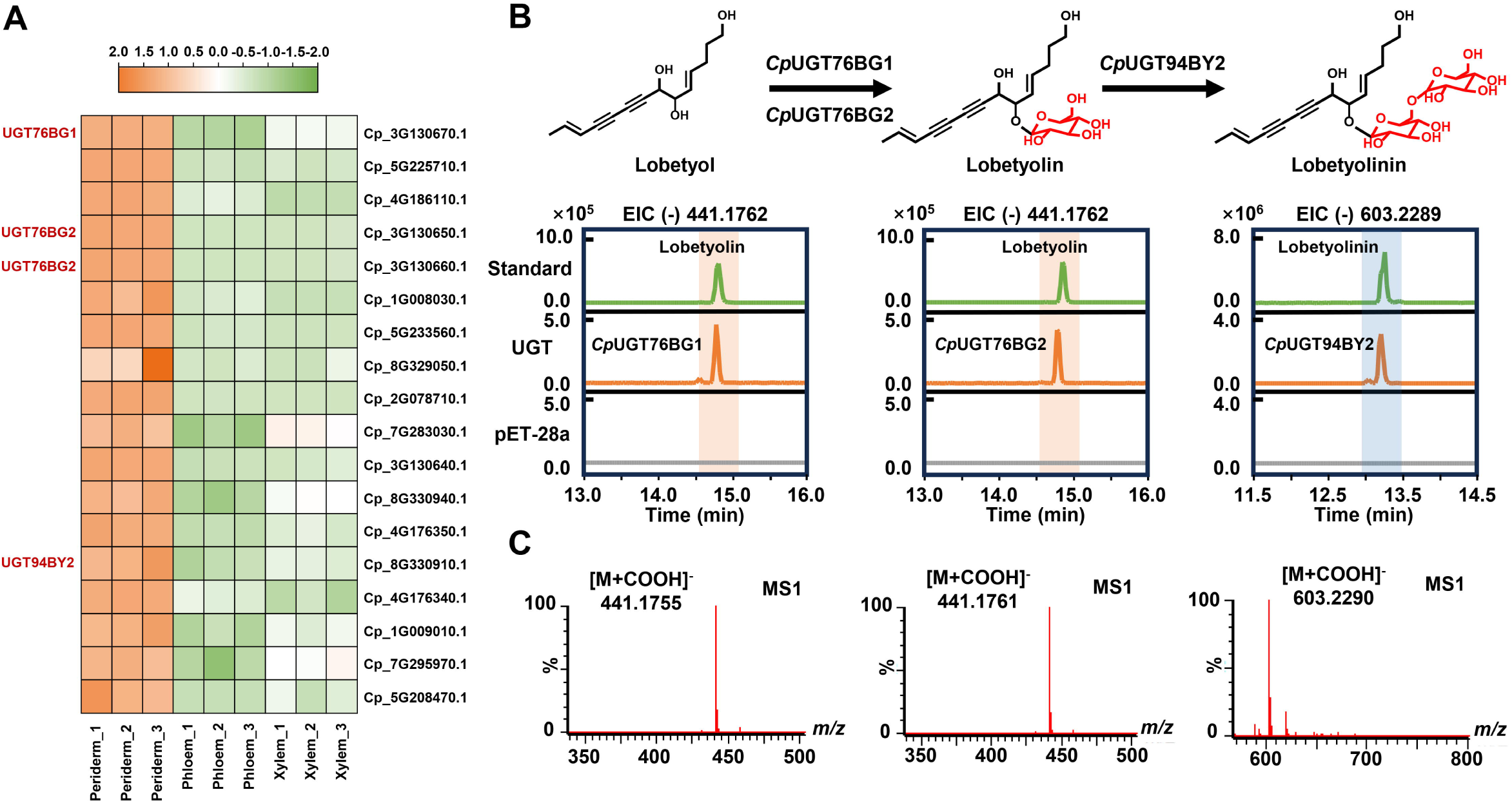
Region-specific screening and in vitro enzymatic assay of UGT genes from *C. pilosula* root. **(A)** Eighteen candidate UGTs showing periderm-enriched expression were identified based on MALDI-MSI spatial metabolite patterns; **(B)** The two-step glycosylation process from lobetyol to lobetyolinin was functionally validated *in vitro*. CpUGT76BG1 and CpUGT76BG2 were confirmed to catalyze the 6-*O*-glycosylation of lobetyol, while CpUGT94BY2 mediated β-1, 6-glycosylation using lobetyolin as substrate. Catalytic functions were assessed via extracted ion chromatogram analysis of reaction products. Assays using lobetyol and lobetyolin as substrates revealed distinct peaks corresponding to lobetyolin and lobetyolinin, respectively, which were authenticated using reference standards. The empty vector (pET-28a) served as a negative control; **(C)** MS/MS spectra further confirmed the identity of the catalytic products, including lobetyolin generated by CpUGT76BG1 and CpUGT76BG2, and lobetyolinin produced by CpUGT94BY2.

*In vitro* functional screening revealed that two UGTs, CpUGT76BG1 and CpUGT76BG2, catalyzed the direct glycosylation of lobetyol to form lobetyolin, identified by its deprotonated molecular ion ([M-H]□: *m/z* 441.175) and a retention time of 14.72 mins (Fig. 3B). A third enzyme, CpUGT94BY2, subsequently converted lobetyolin to lobetyolinin ([M-H]□: *m/z* 603.228; RT = 13.15 min) via β-(1→6) glycosylation (Fig. 3B). The identities of both glycosylated products were confirmed by comparison with authentic standards (Fig. 3B, Supplementary Data 10). The remaining periderm-enriched UGT candidates showed no detectable glycosylation activity toward lobetyol and lobetyolin under standard assay conditions (Supplementary Fig. 26). As a result, this direct glycosylation mechanism with polyacetylene scaffold represents a novel catalytic role for UGT76 family, which were previously associated with steroidal saponin, flavonoids and alkaloids biosynthesis^16, 26^. Similarly, CpUGT94BY2 extends the recognized catalytic scope of di-*O*-glycosyltransferases, which were previously characterized for flavonoid and triterpenoid substrates^27, 28^, to include polyacetylene-derived acceptors. Minor peaks were observed at RT = 14.55 min (preceding the main lobetyolin peak at 14.72 min) in the CpUGT76BG1 and CpUGT76BG2 chromatograms, and at RT = 13.00 min (preceding the main lobetyolinin peak at 13.15 min) in the CpUGT94BY2 chromatogram. High-resolution MS1 and MS2 characterization confirmed that these minor peaks represent structural region-isomers generated by slight *in vitro* regio-promiscuity of the recombinant enzymes under high substrate concentrations. Specifically, the main products (RT = 14.72 min for lobetyolin; RT = 13.15 min for lobetyolinin) exhibit both formate adduct ions and pseudo-molecular ions identical to the authentic standards, whereas the interfering isomers show only the formate adduct ion without the corresponding pseudo-molecular ion due to the lower concentration (Supplementary Figs. 11 and 12).

### *In planta* functional validation of CpUGT76BG1, CpUGT76BG2 and CpUGT94BY2

To verify the *in vivo* functionality of the identified enzymes, we transiently expressed CpUGT76BG1 and CpUGT76BG2 together with lobetyol, and CpUGT94BY2 with lobetyolin, using pHB-based expression vectors in *Nicotiana benthamiana* leaves. LC-MS analysis confirmed lobetyolin production in leaves co-infiltrated with CpUGT76BG1 or CpUGT76BG2 and lobetyol (Fig. 4A). Similarly, CpUGT94BY2 generated lobetyolinin from lobetyolin by comparison with empty vector control group, indicating successful biosynthesis of glycosylation of lobetyol and lobetyolin in *N. benthamiana* (Fig. 4A).

**Figure 4.**
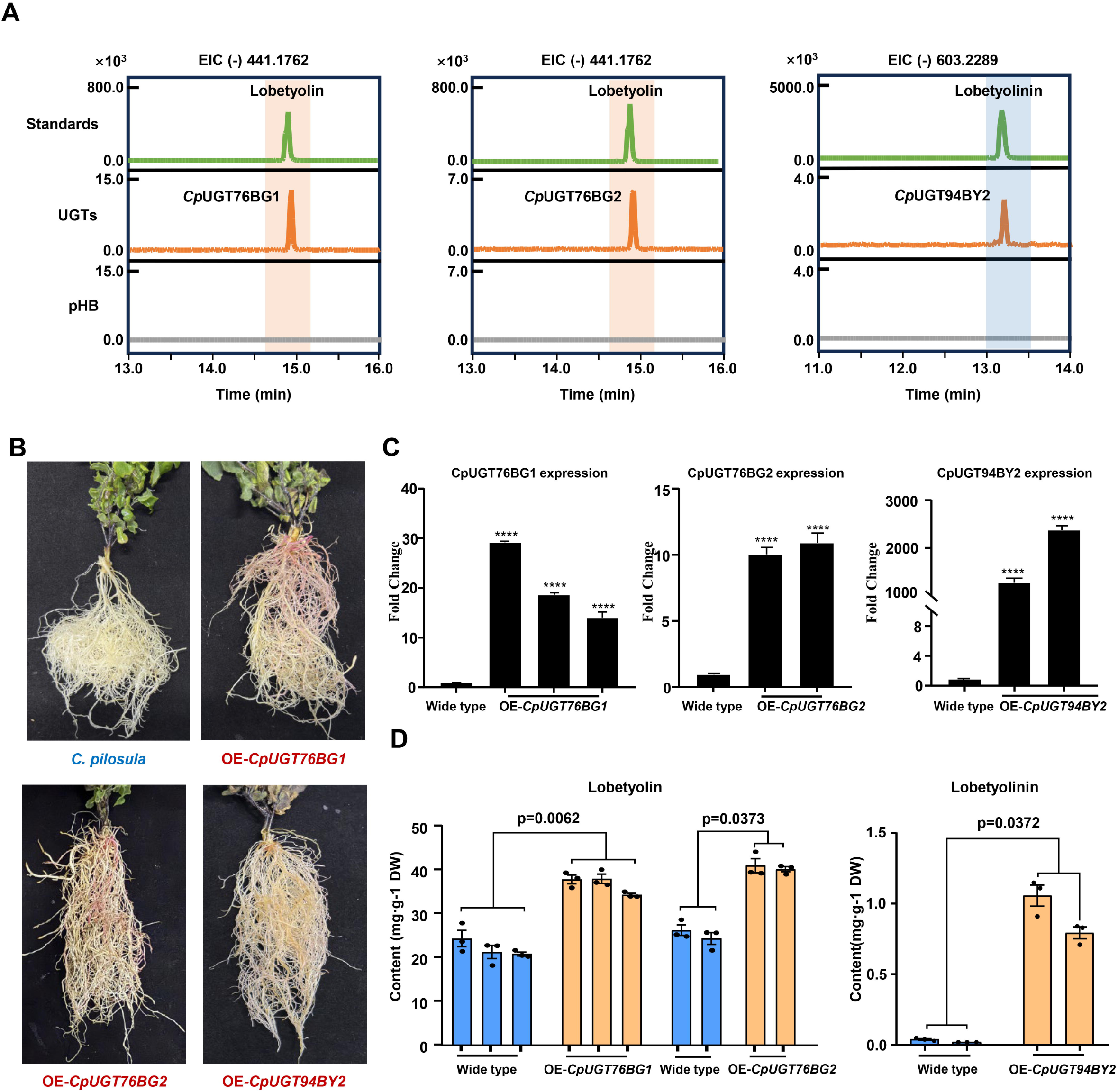
Functional validation of CpUGT76BG1, CpUGT76BG2, and CpUGT94BY2 via transient expression in *N. benthamiana* and stable overexpression *in planta*. **(A)** Transient expression in *N. benthamiana* leaves was performed to assess the catalytic activity of CpUGT76BG1 and CpUGT76BG2 using lobetyol as substrate, and CpUGT94BY2 using lobetyolin. Peaks corresponding to lobetyolin and lobetyolinin were verified with reference standards. The empty vector (pHB) served as a negative control; **(B)** Transgenic *C. pilosula* lines overexpressing each gene are shown, with a scale bar of 1 cm; **(C)** qRT-PCR quantification of transcript levels in the respective transgenic lines. The specific target genes (*CpUGT76BG1*, *CpUGT76BG2*, and *CpUGT94BY2*) are indicated by the titles above each corresponding graph. Data are presented as mean ± standard deviation from three biological replicates; **(D)** Lobetyolin content was measured in roots overexpressing CpUGT76BG1 (OE-CpUGT76BG1) and CpUGT76BG2 (OE-CpUGT76BG2), while lobetyolinin was quantified in roots overexpressing CpUGT94BY2 (OE-CpUGT94BY2). Data are shown as mean ± standard deviation (n = 3). A statistically significant difference in metabolite accumulation between wild-type and transgenic plants was observed, with P < 0.05.

Given the confirmed enzymatic activity established in *E. coli* and *N. benthamiana* systems, we stably transformed *C. pilosula* with constructs encoding CpUGT76BG1, CpUGT76BG2, and CpUGT94BY2 using the pCAMBIA1305-RUBY vector driven by the CaMV 35S promoter (Fig. 4B). Quantitative RT–PCR analysis revealed greater than ten-fold upregulation of the respective transcripts in transgenic roots compared to wild-type plants (Fig. 4C). Metabolite quantification showed that overexpression of CpUGT76BG1 increased lobetyolin accumulation to 36.60 ± 1.40 mg/g dry weight (DW), compared to 22.03 ± 2.22 mg/g DW in wild-type controls. CpUGT76BG2 overexpression elevated lobetyolin to 40.48 ± 1.88 mg/g DW (Fig. 4D, Supplementary Data 14). Furthermore, transgenic lines overexpressing CpUGT94BY2 exhibited an approximately thirtyfold increase in lobetyolinin content, reaching 0.92 ± 0.10 mg/g DW, relative to 0.03 ± 0.01 mg/g in wild-type plants (Fig. 4D, Supplementary Data 14). Given that the CaMV 35S promoter drives ectopic expression across all root tissues, the substantial increases in lobetyolin are likely attributable not only to enhanced enzyme activity in the periderm but also to the utilization of the available lobetyol pool in parenchyma cells, which is uniformly distributed across root tissues as shown by MALDI-MSI (Fig. 2). Our results thus genetically define a two-step glycosylation pathway: CpUGT76BG1 and CpUGT76BG2 catalyze the first glycosylation of lobetyol, and then CpUGT94BY2 acts as a sugar-on-sugar glycosyltransferase to extend the saccharide chain for the production of lobetyolinin.

### Structural basis for substrate recognition and catalytic mechanism

To elucidate the structural basis for polyacetylene glycosylation, we performed amino acid sequence alignments and generated structural models of CpUGT76BG1, CpUGT76BG2, and CpUGT94BY2 using AlphaFold 3.0 and performed molecular docking simulations with lobetyol and lobetyolin as substrates. Structural analysis revealed that CpUGT76BG1 possesses a deep substrate-binding tunnel architecture for lobetyol accommodation (Fig. 5A). The tunnel entry is defined by a constriction ring composed of four residues: Pro83, Asn84, Ile85, and Pro287, with Pro287 positioned as an entrance dome located 8.3 Å from Pro83. The catalytic residue His29 forms a hydrogen bond with the C-14 hydroxyl group of lobetyol, facilitating its deprotonation and subsequent nucleophilic attack on UDP-glucose via an SN2-type mechanism. Asp123 serves as the second catalytic residue within the tunnel (Fig. 5A). The proximal tunnel wall is lined by Val82 and Leu88, while the distal wall comprises Leu125, Leu201, Ile204, and Met208. A distal cap residue, Leu384, is positioned 5.6 Å from the C-3 atom of lobetyol, enclosing the substrate within the tunnel cavity (Fig. 5B).

**Figure 5.**
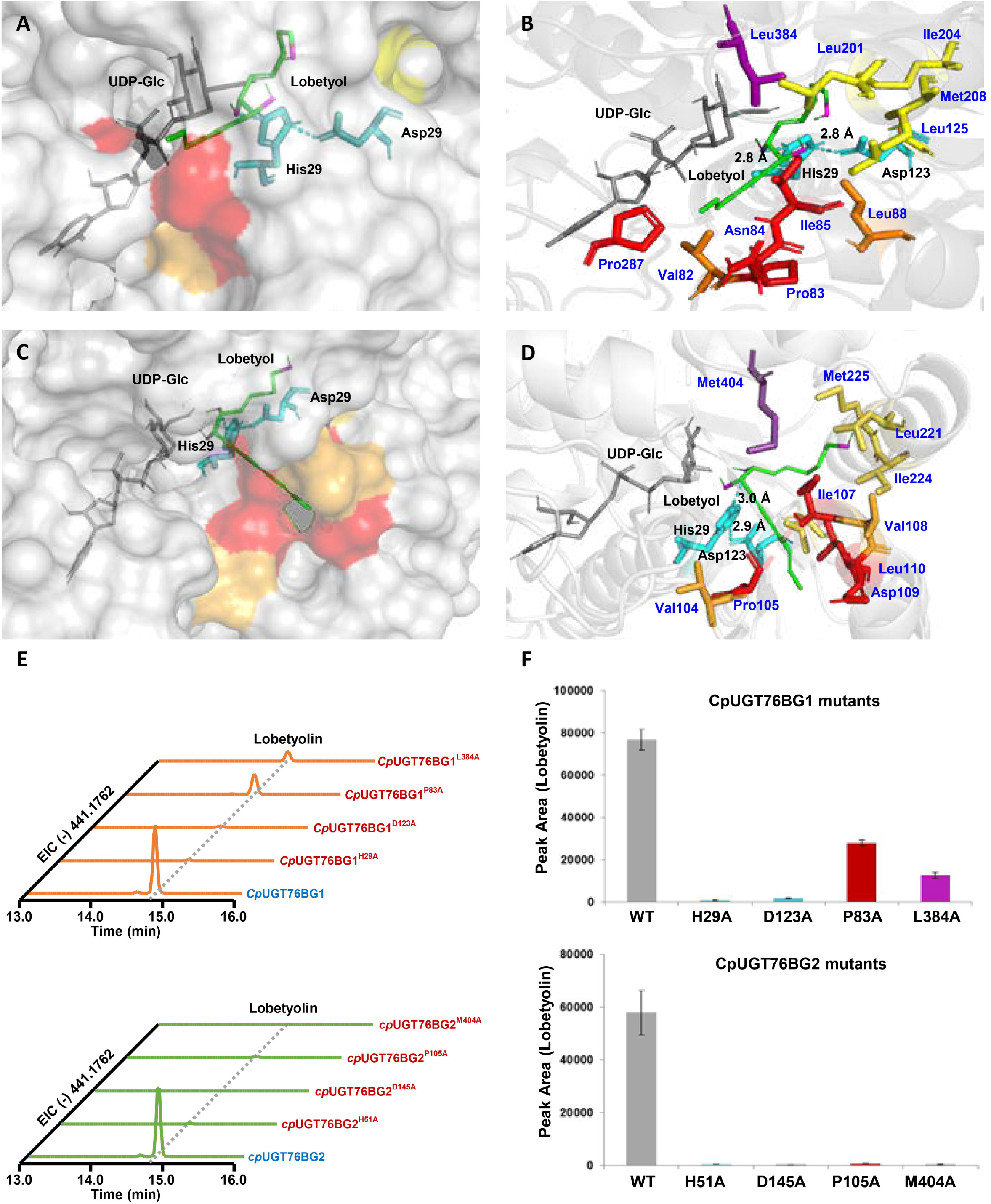
Structural comparison of substrate-binding architecture in CpUGT76BG1 and CpUGT76BG2 and validation by site-directed mutagenesis. **(A)** Surface representation of the AlphaFold3-predicted CpUGT76BG1 model with docked lobetyol (green sticks) and UDP-Glc (gray sticks), showing the deep hydrophobic tunnel with a clipped cross-sectional view. The catalytic dyad (His29, Asp123; cyan) is labeled; **(B)** Cartoon representation of CpUGT76BG1 with individual tunnel residues shown as sticks and color-coded by zonation: entrance constriction ring (Pro83, Asn84, Ile85, Pro287; red), proximal wall (Val82, Leu88; orange), catalytic dyad (His29, Asp123; cyan), distal wall (Leu125, Leu201, Ile204, Met208; yellow), and distal seal Leu384 (purple); **(C)** Surface representation of the AlphaFold3-predicted CpUGT76BG2 model with docked lobetyol (green sticks) and UDP-Glc (gray sticks), showing the deep hydrophobic tunnel with a clipped cross-sectional view. The catalytic dyad (His51, Asp145; cyan) is labeled; **(D)** Cartoon representation of CpUGT76BG2 with individual tunnel residues shown as sticks and color-coded by zonation: entrance constriction ring (Pro105, Ile107, Asp109, Leu110; red), proximal wall (Val104, Val108; orange), catalytic dyad (His51, Asp145; cyan), distal wall (Leu147, Leu221, Ile224, Met225; yellow), and distal seal Met404 (purple); (**E**) Extracted ion chromatograms of wild-type and single-point alanine mutants for both CpUGT76BG1 (H29A, D123A, P83A, L384A) and CpUGT76BG2 (H51A, D145A, P105A, M404A), showing lobetyolin production (*m/z*=441.176); (**F**) Quantification of lobetyolin peak area for wild-type and each mutant of CpUGT76BG1 (above) and CpUGT76BG2 (below).

CpUGT76BG2 similarly encloses lobetyol within a substrate-binding tunnel that fully embeds the ligand, as observed in the solvent-accessible surface model, with the entry delineated by Pro105, Ile107, Asp109, and Leu110 at the C-14 terminus and the catalytic His51 positioned for C-14 hydroxyl deprotonation (Fig. 5C). Met404 was identified as a candidate distal cap residue, structurally analogous to Leu384 in CpUGT76BG1. Despite divergent primary sequences at their binding cavities, the two enzymes share a conserved spatial logic spanning entry, catalytic center, wall lining, and distal cap. For CpUGT94BY2, residues Glu276 and Trp17 were identified as crucial for coordinating the sugar moiety at the C-14 position of lobetyolin to enable β-1, 6-glycosylation, while Lys96 appeared to stabilize the substrate-binding pocket through a hydrogen bond with the C-1 hydroxyl group (Supplementary Fig. 19). Site-directed mutagenesis employing single-residue alanine substitutions was performed to systematically validate the predicted tunnel architecture. For CpUGT76BG1, mutations targeting each functional zone—catalytic dyad (H29A, D123A), entrance constriction (P83A), and distal seal (L384A)—were generated and assayed *in vitro* (Fig. 5E). The catalytic mutations H29A and D123A reduced activity by approximately 89-fold and 46-fold, respectively, confirming their essential roles in substrate activation (Fig. 5F). In contrast, the structural mutations P83A and L384A resulted in more moderate reductions of approximately 2.7-fold and 6.1-fold, respectively (Fig. 5F), indicating that the entrance constriction and distal seal contribute to but are not individually essential for catalytic competence in CpUGT76BG1. Strikingly, the equivalent mutations in CpUGT76BG2—H51A, D145A, P105A, and M404A—produced a markedly different profile (Fig. 5G). While the catalytic mutations H51A and D145A abolished activity by approximately 132-fold and 250-fold, respectively (Fig. 5H), the structural mutations P105A and M404A also nearly eliminated activity, with reductions of approximately 83-fold and 186-fold (Fig. 5H). This pronounced asymmetry, where structural tunnel mutations cause only 2.7- to 6.1-fold reduction in CpUGT76BG1 but 83- to 186-fold reduction in CpUGT76BG2, reveals fundamentally different degrees of architectural robustness between the two paralogs. For CpUGT94BY2, mutations at Trp17 and Glu276 completely abrogated glycosylation activity, while the K96A variant severely disrupted substrate binding, highlighting the functional importance of hydrogen bonding in both sugar transfer and transition-state stabilization (Supplementary Fig. 19).

### Substrate and sugar donor specificity of CpUGT76BG enzymes

To further define the substrate and sugar donor specificity of CpUGT76BG1 and CpUGT76BG2, we challenged the enzymes with structurally related polyacetylene analogs and alternative nucleotide sugar donors. When tested against falcarindiol (a C17 natural polyacetylene diol structurally related to lobetyol)^29^, CpUGT76BG1 exhibited detectable glycosylation activity, whereas CpUGT76BG2 did not (Supplementary Fig. 22). In contrast, neither enzyme catalyzed the glycosylation of 10-undecyn-1-ol (a C11 simple alkynol) or 4-pentyn-1-ol (a C5 simple alkynol).

Furthermore, when alternative nucleotide sugar donors (UDP-Galactose, UDP-Glucuronic acid, UDP-Rhamnose, and UDP-Xylose) were tested in place of UDP-Glucose, neither enzyme accepted UDP-Galactose or UDP-Glucuronic acid, but both CpUGT76BG1 and CpUGT76BG2 produced detectable product peaks corresponding to xylosylated and rhamnosylated derivatives of lobetyol (formate adduct ions *m/z* 411.1655 and 425.1810, respectively), indicating that while UDP-Glucose is the preferred sugar donor, the UGT76BG tunnel architecture can accommodate UDP-Xylose and UDP-Rhamnose as alternative donors. CpUGT94BY2 similarly accepted UDP-Xylose and UDP-Rhamnose, generating the corresponding lobetyolinin derivatives (formate adduct ions *m/z* 573.2179 and 587.2334, respectively (Supplementary Fig. 23). To quantify the catalytic efficiency of each UGT, we performed steady-state kinetic analysis under saturating UDP-Glc conditions with varying acceptor substrate concentrations (Supplementary Fig. 13). CpUGT76BG1 catalyzed lobetyol glycosylation with a *K_m_* of 15.1 μM, a *k_cat_*of 0.050 s□¹, and a catalytic efficiency (*k_cat_/K_m_*) of 0.003 μM□¹·s□¹. CpUGT76BG2 exhibited a higher *K_m_* (27.6 μM), a lower *k_cat_* (0.028 s□¹), and a 3.3-fold lower *k_cat_/K_m_* (0.001 μM□¹·s□¹). CpUGT94BY2, operating on lobetyolin, displayed the highest catalytic efficiency at 0.012 μM□¹·s□¹ (*K_m_* = 11.3 μM; *k_cat_* = 0.142 s□¹). The resulting hierarchy is consistent with the structural model in which robust tunnel of CpUGT76BG1 sustains higher turnover than fragile tunnel of CpUGT76BG2, while CpUGT94BY2 achieves the highest efficiency on a pre-glycosylated substrate with greater aqueous accessibility.

### Evolutionary innovation of the UGT76 family in polyacetylene glycosylation

Campanulaceae species consistently accumulate polyacetylene glycosides such as lobetyolin, suggesting a lineage-specific metabolic innovation within this plant family. LC–MS analysis confirmed the presence of lobetyol and its glycosylated derivatives, lobetyolin and lobetyolinin, in both root and leaf tissues of four Campanulaceae species: *P. grandiflorus*, *Codonopsis tubulosa*, *Pseudocodon convolvulaceus*, and *C. lanceolata* (Supplementary Fig. 24). To elucidate the genomic basis of this chemical trait and the evolutionary history of the UGT76 family, we examined the phylogenomic and synteny analyses containing *UGT76BG1*, *UGT76BG2*, and related *UGT76* genes across Campanulaceae species (*C. pilosula*, *P. grandiflorus*, *L. seguinii*, and *C. lanceolata*) and outgroup taxa (*Aralia elata*, *Scutellaria baicalensis*, and *Vitis vinifera*) (Fig. 6A). All species included Campanulaceae famly only share core eudicot γ WGD event, except for *L. seguinii* exhibited an additional recent WGD (*K*s ≈ 0.1) (Fig. 6A). Synteny comparison with ancestral eudicot model of *V. vinifera* revealed a conserved genomic block predating the divergence of Campanulaceae (Fig. 6A). Distinctive evolutionary profile of UGT76s have shown that lineage-specific tandem duplications have expanded the copy number exclusively within Campanulaceae, with more than six *UGT76* genes retained in the syntenic segment (Fig. 6A); flanking non-UGT genes outside the UGT76 locus remain deeply conserved across Campanulaceae species across an extended region encompassing 15 additional genes on each side of the cluster, confirming that the tandem expansion is strictly confined to the UGT76 genes alone (Supplementary Fig. 25, Supplementary Data 16). Systematic functional analysis of the collinear UGT76 loci across Campanulaceae species further established the prevalence of lobetyol glycosylation activity among tandemly duplicated paralogs. Except for *UGT76BG1* and *UGT76BG2*, we chemically synthesized the full-length coding sequences of 8 selected tandemly duplicated UGT76 orthologs from the distinct Campanulaceae species and expressed them as recombinant proteins in *E. coli* to test their catalytic activity against lobetyol *in vitro*. We confirmed that the functional activities were found in two *UGT76* genes from *P. grandiflorus* (Pg_chr01_16800T and Pg_chr01_16810T), four from *L. seguinii* (Ls_Gene005190.t01, Ls_Gene005191.t01, Ls_Gene006333.t01, and Ls_Gene006334.t01), and two from *C. lanceolata* (Cl_chr03_42420T and Cl_chr03_42430T) (Fig. 6B). In total, eleven orthologs arising from lineage-specific tandem duplications, including the 8 tested here and the 3 from *C. pilosula*, catalyzed the conversion of lobetyol to lobetyolin (Fig. 6B). Strikingly, five UGT76 genes within the Campanulaceae branch that lack collinear relationships with the conserved syntenic block showed no detectable lobetyol glycosylation activity (Fig. 7, Supplementary Fig. 26), establishing a strict correlation between synteny conservation and functional retention: all collinear UGT76 paralogs catalyze lobetyol, whereas all non-collinear ones lack this activity. These findings indicate that neofunctionalization driven by tandem duplication, constrained within the syntenic genomic context, enabled the adaptation of UGT76 enzymes to linear polyacetylene substrates.

**Figure 6.**
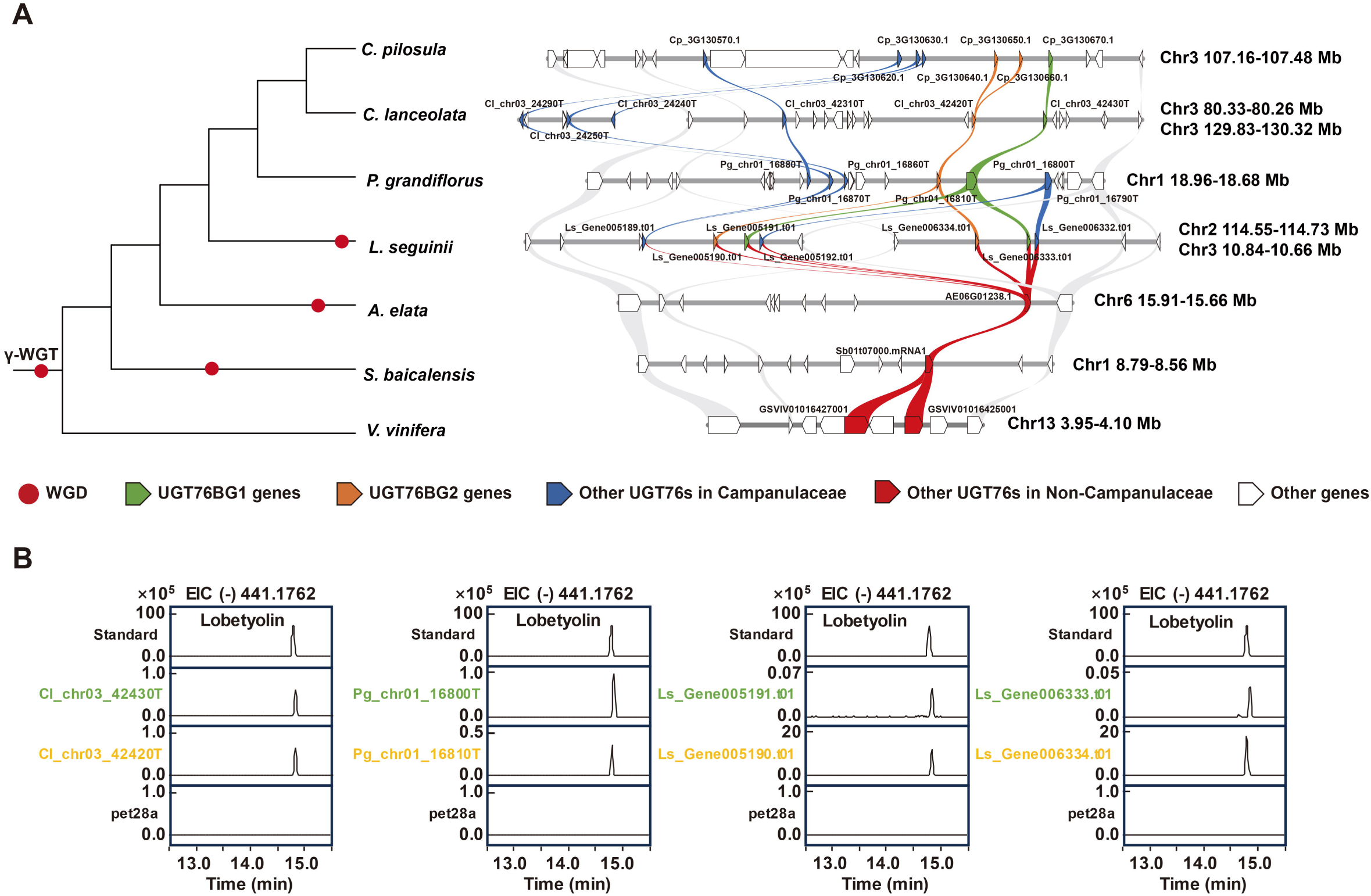
Phylogenomic and syntenic analyses reveal tandem duplication-driven evolution of the UGT76 family in Campanulaceae. **(A)** Phylogenomic divergence of *C. pilosula* compared to three other Campanulaceae species (*L. seguinii*, *P. grandiflorus*, and *C. lanceolata*), alongside the syntenic analysis of genomic regions containing UGT76 orthologs. Red circle indicates the relative timing of the WGD event. Green and orange ribbons denote syntenic UGT76BG1 and UGT76BG2 genes that accepted lobetyol as substrate, whereas blue ribbons denote syntenic UGT76s lacking lobetyol glycosylation activity; **(B)** *In vitro* functional characterization of collinear UGT76 enzymes from *L. seguinii*, *P. grandiflorus*, and *C. lanceolata*. The empty vector (pET-28a) served as a negative control. A peak corresponding to lobetyolin was verified with reference standards.

**Figure 7.**
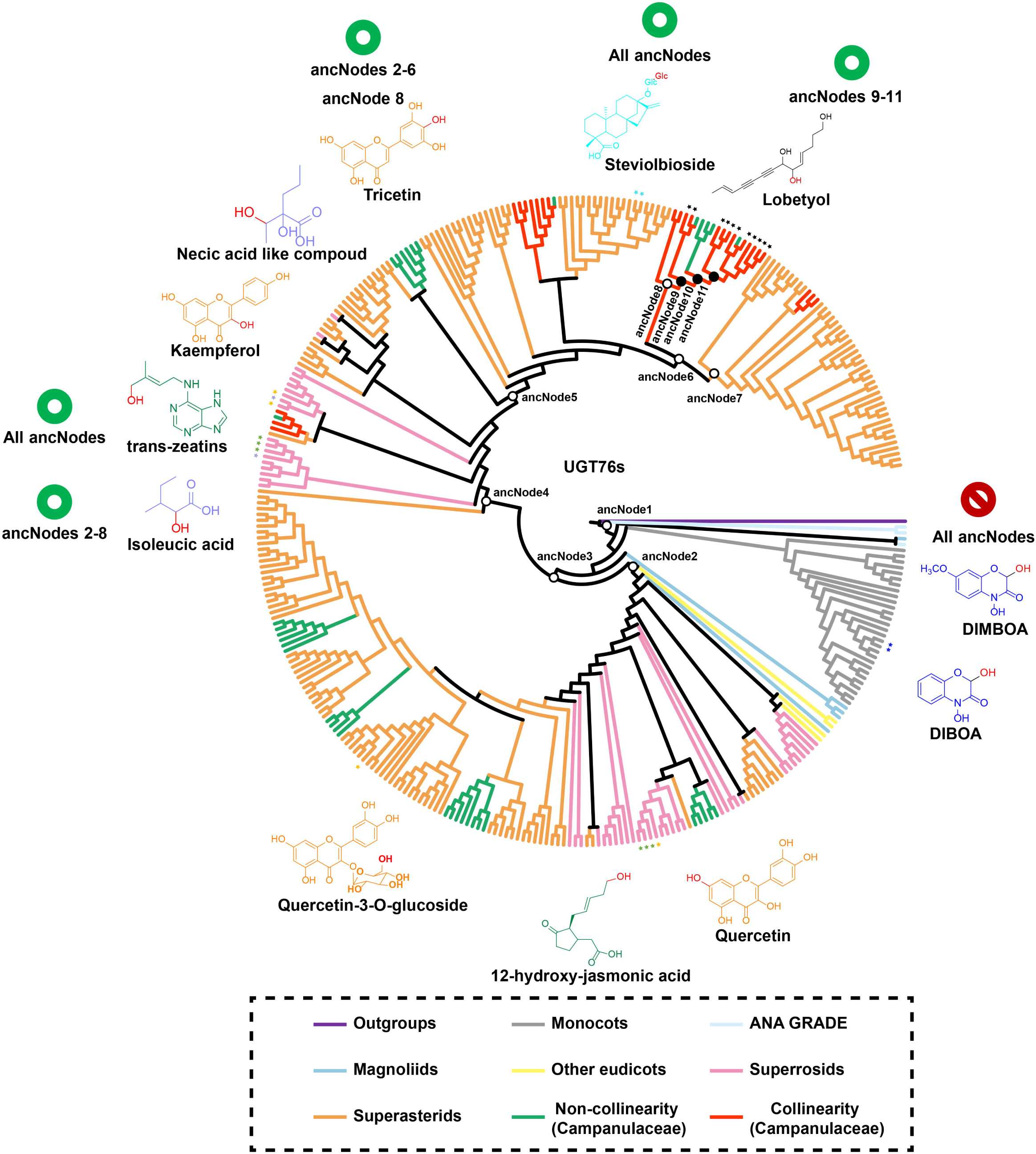
Evolutionary history of the UGT76 family across angiosperms. A phylogenetic tree illustrates the diversification of the UGT76 family based on a broad sampling of angiosperm species. Eleven ancestral nodes were reconstructed and functionally characterized to elucidate evolutionary transitions within this enzyme family. Black solid ancestral nodes have the function of catalyzing lobetyol, while the black hollow nodes cannot accept it. Within the Campanulaceae branch, eleven collinear UGT76 genes (red bar) form a monophyletic clade with lobetyol glycosylation activity, whereas five non-collinear UGT76 genes (green bar) lack this catalytic function. The corresponding functional genes are labeled with star-shaped markers. The phylogram is color-coded according to major plant lineages, including ANA Grade, monocots, magnoliids, other eudicots, superrosids, and superasterids. Chemical structures surrounding the tree represent accepted substrates of functionally characterized UGT76 enzymes, encompassing fatty acid derivatives, flavonoids, phytohormones, alkaloids, terpenoids, and polyacetylenes. Among these, six substrates (trans-zeatin, steviolbioside, tricetin, isoleucic acid, DIMBOA, and lobetyol) were experimentally tested in enzyme activity assays with the reconstructed ancestral enzymes. Green circles next to each substrate indicate that the corresponding ancestral enzyme can catalyze that substrate, whereas red prohibition circles indicate no detectable catalytic activity. Detailed annotations of the phylogeny are provided in Supplementary Fig. 27.

To trace the evolutionary origin and substrate specificity trajectory of this catalytic function, we reconstructed a phylogeny of UGT76 enzymes from diverse angiosperms and performed ancestral sequence resurrection for eleven key nodes (AncNode-1 through AncNode-11) (Fig. 7, Supplementary Fig. 27). Substrates accepted by UGT76s, as illustrated around the tree, include fatty acid derivatives, flavonoids, phytohormones, alkaloids, and terpenoids (Fig. 7).To determine the substrate specificity trajectory underlying polyacetylene glycosylation innovation, we performed comprehensive activity profiling of eleven resurrected ancestral enzymes and the two extant enzymes (CpUGT76BG1 and CpUGT76BG2) against five representative substrates spanning the known UGT76 family repertoire: trans-zeatin (cytokinin), steviolbioside (diterpene glycoside), tricetin (flavonoid), isoleucic acid (amino acid derivative), and DIMBOA (hydroxamic acid), in addition to lobetyol (polyacetylene aglycone) (Fig. 7). The deepest ancestor (AncNode-1) exhibited robust activity toward trans-zeatin and steviolbioside, but no detectable activity toward tricetin, isoleucic acid, DIMBOA, or lobetyol, indicating that the earliest ancestor retained only the most conserved UGT76 functions. Notably, trans-zeatin and steviolbioside were catalyzed by all 13 enzymes (AncNode-1 through AncNode-11, along with CpUGT76BG1 and CpUGT76BG2), establishing these two substrates as the most ancient and evolutionarily stable UGT76 activities. In contrast, the ability to catalyze DIMBOA emerged only in earlier-diverging sublineages descending from AncNode-1, and was not present in the deepest ancestor itself, indicating that DIMBOA catalysis was a later acquisition in specific UGT76 subfamilies. From AncNode-2 to AncNode-6 and AncNode-8, ancestral enzymes acquired broad-spectrum activity toward both tricetin and isoleucic acid while retaining trans-zeatin and steviolbioside activity, representing a functional expansion that coincides with the shallow-pocket architecture. A notable exception is AncNode-7, a parallel lineage branching from AncNode-6 that does not lead to extant CpUGT76BG1/BG2; AncNode-7 retains isoleucic acid activity but lacks both tricetin and lobetyol activities, demonstrating that tricetin and isoleucic acid recognition are decoupled and controlled by distinct residue combinations. Starting precisely at AncNode-9, a profound functional switch occurred: activity toward tricetin and isoleucic acid was completely abolished, while robust lobetyol glycosylation activity emerged for the first time. AncNode-10 and AncNode-11, as well as the extant CpUGT76BG1/BG2, all retained this same activity profile (trans-zeatin, steviolbioside, lobetyol), confirming that polyacetylene specialization is a derived, Campanulaceae-specific innovation (Fig. 7, Supplementary Fig. 29).

## 4. Discussion

The integration of a T2T-level genome with spatial metabolomics and ancestral sequence resurrection provides a molecular insights into how lineage-specific metabolic innovations emerge in plant specialized metabolism. Our study establishes not only the genetic and biochemical basis of polyacetylene glycoside biosynthesis in Campanulaceae, but also reveals a complete evolutionary trajectory of substrate specificity within the UGT76 family—from conserved hormonal glycosylation to defensive polyacetylene specialization. This trajectory exemplifies how structural innovation drives functional divergence, a principle that extends far beyond the UGT family.

This shared spatial enrichment in the periderm provided a strategic focus for simultaneously identifying genes from multiple pathway steps, demonstrating the efficiency and utility of spatial heterogeneity-guided gene discovery. It is noteworthy that imposing an additional filter for xylem-high expression would have inadvertently excluded UGT76BG2, underscoring the practical advantage of prioritizing the most pronounced spatial commonality of periderm enrichment as a singular criterion for candidate selection. This imaging-guided approach enabled precise identification of the UGT76 and UGT94 families, circumventing resource-intensive candidate screening. The methodology holds substantial promise for elucidating other spatially compartmentalized pathways, performing at the level of single cells and high-resolution determination of gene regulatory features for solving the monoterpene indole alkaloid biosynthesis in *Catharanthus roseus*^30^. To contextualize these glycosylation events within the broader polyacetylene biosynthetic framework, we proposed a de novo pathway from stearic acid (18:0) to lobetyolinin (Supplementary Fig. 30). The committed entry into polyacetylene metabolism is catalyzed by divergent FAD2 variants with Δ12-acetylenase activity^31, 32^. Downstream steps—peroxisomal β-oxidation (ACX/MFP/KAT) for C18 → C14 chain truncation, allylic isomerization, and CYP450-mediated hydroxylation—remain proposed but unverified^33–36^, representing key targets for future characterization. The identification of UGT76BG1/BG2 and UGT94BY2 now provides the terminal glycosylation steps of this pathway, linking the polyacetylene scaffold generation to the accumulation of bioactive glycosides (Supplementary Fig. 30).

The catalytic mechanism of CpUGT76BG1 and CpUGT76BG2 is canonical: both employ the conserved His-Asp dyad (His29/Asp123; His51/Asp145) that characterizes virtually all Plant Family 1 UGTs. The novelty of these enzymes therefore resides not in catalytic chemistry but in the structural strategy for substrate positioning. The most striking discovery of our study is that CpUGT76BG1 and CpUGT76BG2 have evolved a deep hydrophobic tunnel to accommodate linear, highly hydrophobic polyacetylene scaffolds, representing a structural adaptation unprecedented among characterized plant UGTs. This represents a profound divergence from the canonical UGT architecture, which predominantly employs a shallow, open binding cavity where planar aromatic substrates (such as flavonoids or phenylpropanoids) are stabilized by π–π stacking interactions within the binding cavity^37, 38^.

Polyacetylene substrates present a fundamentally different challenge: their rigid carbon–carbon triple bonds resist conformational bending, and their complete lack of polar functional groups eliminates conventional hydrogen-bonding anchors. The tunnel architecture is not a simple “deepening” of the shallow pocket, but a coordinated multi-residue innovation that creates a continuous hydrophobic environment along the entire length of the polyacetylene chain. Comparison with the closest structurally characterized paralog—SrUGT76G1 (PDB: 6KVK) from *Stevia rebaudiana*^39^, which possesses a shallow bowl-shaped pocket for diterpene glycoside recognition—reveals the extent of this structural divergence. In SrUGT76G1, the entrance region is guarded by polar residues (Thr285, Ser284) that coordinate sugar moieties near the protein surface. In CpUGT76BG1, these corresponding positions have been repurposed into a constricted, hydrophobic tunnel gateway (Pro83, Asn84, Ile85, Pro287) that seals the substrate inside (Fig. 5B). The substrate specificity analysis further reveals that the tunnel architecture imposes stringent positional constraints on hydroxyl acceptor groups: neither CpUGT76BG1 nor CpUGT76BG2 could glycosylate simple alkynols (10-undecyn-1-ol, 4-pentyn-1-ol) bearing only a terminal primary hydroxyl, in contrast to lobetyol and falcarindiol, which possess an allylic hydroxyl group along the polyacetylene chain. This inability indicates that a terminal primary hydroxyl is not a competent acceptor for the UGT76BG tunnel architecture, consistent with the structural model in which the deep hydrophobic tunnel positions the substrate such that only an internal allylic hydroxyl can be properly oriented toward the catalytic histidine. The mutagenesis data uncover a striking asymmetry in tunnel robustness between the two paralogs: structural mutations at the entrance and distal seal cause only modest activity loss in CpUGT76BG1 but near-complete ablation in CpUGT76BG2 (Figs. 5E-5H). This contrast reveals that the CpUGT76BG1 tunnel retains substrate-positioning capacity even when individual structural constraints are compromised, whereas the CpUGT76BG2 tunnel collapses upon disruption of any single component. This differential robustness mechanistically explains the observed substrate selectivity divergence. The structurally robust tunnel of CpUGT76BG1 accommodates broader polyacetylene substrates including falcarindiol (Supplementary Fig. 22), whereas the structurally fragile tunnel of CpUGT76BG2 enforces stringent specificity, offering a coherent rationale for the evolutionary retention of both paralogs despite their apparent functional redundancy.

Ancestral sequence resurrection across 11 phylogenetic nodes reveals a fundamental principle of UGT76 evolution: the coexistence of evolutionarily stable conserved activities and a highly evolvable substrate-recognition periphery. Importantly, ancestral substrate profiling reveals that this structural innovation, namely the deep hydrophobic tunnel, is strictly correlated with a three-stage functional trajectory: from conserved hormone/diterpene glycosylation (Stage I, AncNode-1), to broader secondary metabolite recognition (Stage II, AncNode-2 through AncNode-8, except AncNode-7), and finally to polyacetylene specialization (Stage III, AncNode-9-11) (Fig. 7, Supplementary Fig. 29). The universal retention of trans-zeatin and steviolbioside glycosylation across all reconstructed enzymes establishes these two substrates as the deepest, most conserved UGT76 activities—a functional core that provides a stable catalytic scaffold tolerating extensive binding-pocket remodeling without catastrophic loss of essential hormone/diterpene glycosylation. Crucially, the transition from the shallow-pocket architecture to the deep-tunnel architecture represents a discrete, coordinated multi-residue substitution event rather than gradual drift. Four co-segregating residue changes at the AncNode-8 → 9 transition — Ser → Pro at the entrance (constricting the gateway), Val→Leu at the proximal wall (expanding the hydrophobic sheath), HINPMLQL→HISPMLHL in the catalytic microenvironment (restoring the hydrogen-bond network), and CFGD/YFGD→GILD at the C-terminal seal (replacing a polar cap with a hydrophobic one), which collectively transform an open, promiscuous binding cavity into a narrow, specialized tunnel. The fact that AncNode-7 (a parallel lineage that retains the shallow-pocket architecture) cannot glycosylate lobetyol despite possessing broad-spectrum activity toward flavonoids and amino acid derivatives proves that the deep tunnel is a necessary condition for polyacetylene recognition, not merely a facilitative adaptation. This necessity suggests that metabolic novelty in plants often requires coordinated multi-residue substitutions that collectively create a new adaptive peak, rather than a single rate-limiting mutation as documented in classical models of enzyme evolution.

The CP2.0 T2T genome assembly was instrumental in revealing the genomic architecture underlying this evolutionary innovation. Contig-based assemblies cannot resolve gene clusters embedded in repeat-dense centromeric regions; the 270 Mb of newly assembled sequence in CP2.0 (Figs. 1A–C, Supplementary Data 6-8) was therefore essential for capturing the full complement of tandemly duplicated UGT76 genes, including two identical CpUGT76BG2 paralogs generated by the most recent tandem duplication that were collapsed into a single annotation in the previous assembly. This genomic expansion provided the genetic raw material for metabolic innovation, mirrors the neofunctionalization paradigm documented in morphine biosynthesis (codeinone reductase (COR)/codeine *O*-demethylase (CODM)/thebaine 6-*O*-demethylase (T6ODM) in Papaveraceae) and flavonoid diversification (CYP82D in *Scutellaria*)^40, 41^. Lineage-specific tandem duplications expanded the UGT76 copy number exclusively within Campanulaceae, with more than six UGT76 genes retained in the syntenic segment (Fig. 6A). The redundancy of CpUGT76BG1 and CpUGT76BG2, conserved across Campanulaceae species, underscores the selective importance of maintaining robust polyacetylene glycoside biosynthesis—likely driven by ecological pressures such as defense against soil-borne pathogens and herbivores^36^.

Several limitations should be noted. First, the structural models presented here are based on AlphaFold3 predictions. Second, loss-of-function genetic validation in *C. pilosula* is confounded by the extensive tandem duplication-driven redundancy of UGT76BG paralogs, and will require multiplex CRISPR approaches capable of simultaneously disrupting the entire paralog cluster. Third, the upstream polyacetylene scaffold-generating steps, including FAD2 acetylenase, β-oxidation, and CYP450-mediated hydroxylation, remain uncharacterized and represent the next priorities for completing the biosynthetic pathway from primary metabolism to bioactive glycosides. Beyond these limitations, the application of these deep-tunnel UGTs in synthetic biology platforms offers promising avenues for the sustainable production of bioactive polyacetylene glycosides; in particular, CpUGT76BG1, which glycosylates a broader range of polyacetylene chain substrates including lobetyol and falcarindiol, represents a versatile catalytic element for polyacetylene glycoside engineering. The well-defined substrate specificity and preferential sugar donor selectivity delineated here, where UDP-Glucose serves as the primary donor while UDP-Xylose and UDP-Rhamnose are tolerated as alternatives (consistent with their shared pyranose scaffold geometry) and UDP-Galactose and UDP-Glucuronic acid are rejected, provides a defined structural template for rational enzyme engineering.

In summary, this study demonstrates how the integration of spatial metabolomics, high-quality genomics, and evolutionary analysis can uncover deep-rooted co-evolutionary trajectories between genes and metabolites in natural systems. The discovery of a tunnel-based substrate envelopment strategy specifically adapted for linear polyacetylene chains expands the recognized structural repertoire of plant UGTs, while the ancestral reconstruction of a stepwise remodeling trajectory provides a molecularly detailed account of how tandem duplication-driven neofunctionalization generates metabolic novelty. This multimodal approach not only identifies novel biosynthetic enzymes with unique substrate specificities but also provides molecular insights into how tandem duplication drives targeted neofunctionalization in plant specialized metabolism.

## Materials and Methods

### Plant Materials

Fresh *Codonopsis pilosula* specimens were collected in Wen County, Longnan City, Gansu Province (September 2023). Periderm, xylem, and phloem tissues from *C. pilosula* roots were dissected for RNA-Seq (n=3) and metabolite analysis (n=6). Campanulaceae species, *Platycodon grandiflorus*, *Codonopsis tubulosa*, *Pseudocodon convolvulaceus and Codonopsis lanceolata*, were collected in Jilin Agricultural University and authenticated by Professor Peng Di (Research Center of Ginseng Breeding and Application, Jilin Agricultural University). Voucher specimens are archived at the Institute of Chinese Materia Medica, Shanghai University of Traditional Chinese Medicine.

### Chemical standards and sample preparation

#### Reagents were sourced as follows

HPLC-grade formic acid (Fisher Scientific, Fair Lawn, NJ, USA); distilled water (Watsons, Hong Kong, China); warfarin internal standard (Sigma-Aldrich, Madrid, Spain); leucine enkephalin (Waters, Milford, MA, USA); HPLC-grade acetonitrile and methanol (Merck, Darmstadt, Germany); and 17 chemical reference standards (>98% purity, Shanghai Standard Technology, Shanghai, China). Structural details for standards are provided in Supplementary Data 10. The structures of compounds lobetyol (aglycone), lobetyolin (monoglycoside), lobetyolinin (diglycoside), and pratianlin B (triglycoside) were confirmed by nuclear magnetic resonance (NMR, 400 MHz) spectroscopy by both 1D ^1^H-NMR and ^13^C-NMR (Supplementary Figs. 15-18 and Supplementary Data 18-19)^42, 43^.

Root and leaf samples were freeze-dried (Scientz-100F), pulverized to homogeneous powders, and sieved (40-mesh). Powders (0.01 g) underwent ultrasonication (53 kHz, 350 W, 60 min, RT) in 1 mL 70% methanol (*v*/*v*) containing warfarin (5 μg/mL). After centrifugation (12,000 × *g*, RT), supernatants were stored at 4°C pending UPLC-QTOF-MS analysis. A quality-control sample was generated by combining 10 μL aliquots from each test sample.

For MALDI-MSI, *C. pilosula* roots were sectioned (2-4 cm), embedded in 5% carboxymethyl cellulose (CMC), and cryofixed for 5 min after solidification. Cryosections (50 μm thickness) were obtained using a microtome (blade advancement: 10-100 μm), mounted on glass slides, labeled, and cryopreserved. Sections were air-dried (RT) prior to analysis. Optical microscopy imaging preceded MALDI-MSI processing.

#### LC-MS and MALDI-MSI-based metabolite profiling

Chromatographic separation was performed using an ACQUITY UPLC system (Waters, Milford, MA, USA) equipped with a binary pump, autosampler, and column compartment. Separation occurred on an ACQUITY UPLC T3 column (2.1 × 100 mm, 1.8 μm) maintained at 40°C, with mobile phases: (A) 0.1% (*v*/*v*) aqueous formic acid and (B) 0.1% (*v*/*v*) formic acid in acetonitrile. The gradient elution program was: 2-5% B (0-6 min), 5-15% B (6-10 min), 15-20% B (10-13 min), 20-22% B (13-16 min), 22-60% B (16-20 min), 60-75% B (20-24 min), 75-95% B (24-30 min), isocratic 95% B (30-33 min), followed by re-equilibration at 2% B (33-36 min). The injection volume was 2 μL with a flow rate of 0.40 mL/min.

Mass detection employed a Xevo G2-XS QTOF mass spectrometer (Waters, Milford, MA, USA) with electrospray ionization under these parameters: sample cone 30 V, source temperature 150°C, desolvation temperature 450°C, cone gas flow 50 L/h, desolvation gas flow 600 L/h, capillary voltage 2.5 kV (negative mode) or 3 kV (positive mode), mass range 50-1200 m/z, scan time 0.35 s, low collision energy 6 eV (negative)/10 eV (positive), high collision energy 30 eV. Leucine enkephalin (2 ng/mL) provided lock mass correction, while sodium formate (0.5 mM) enabled instrument calibration. Data processing used MassLynx v4.2 (Waters, Milford, MA, USA).

For MALDI-MSI, tissue sections mounted on sample stages underwent sequential optimization: Z-axis focusing for image clarity, X/Y-axis positioning for region selection, and final Z-axis adjustment to minimize laser spot size while maximizing signal intensity. Analyses used a TransMIT AP-SMALDI 5AF ion source coupled to a Thermo Scientific Orbitrap Exploris 120 mass spectrometer in positive-ion mode. Parameters included: mass range 70-1050 m/z, spatial resolution 60 μm. Matrix DHB (30 mg/mL in methanol: water 70:30 *v*/*v*) was applied at 20 μL/min flow rate under N□gas (5 L/min), with a nozzle speed of 8 mm/s and duration of 50 min/slide. Pre-deposition rinsing used methanol:water (70: 30 *v*/*v*) at 30 μL/min with identical N□flow for 5 min.

#### Genome sequencing, assembly, and annotation

High-quality genomic DNA was extracted from fresh leaves of a single *C. pilosula* plant and used to construct multiple sequencing libraries. Sequencing was performed using a combination of short-read and long-read technologies. Short-insert libraries were prepared and sequenced on the DNBSEQ-T7 platform (MGI, Anoroad, Beijing, China). For long-read sequencing, libraries were constructed and sequenced both on the PromethION platform (Oxford Nanopore Technologies, Oxford, UK) for ultra-long reads and on the PacBio Sequel II system (PacBio, Menlo Park, CA, USA) for high-fidelity (HiFi) reads. Additionally, a Hi-C library was prepared from young leaves of *C. pilosula* according to a standard protocol and subjected to paired-end sequencing ^44^.

Genome size and heterozygosity were estimated from DNBSEQ-T7 data using Jellyfish v2.2.10^45^ and GenomeScope v2.0^46^ under default parameters. Primary contig assembly was performed with hifiasm (v0.20.0)^47^ using HiFi, Hi-C, and ONT reads. The resulting contigs were subsequently processed with Purge_dups (v1.2.1) ^48^ to remove redundant overlaps. To improve assembly quality, long-read assembly was further conducted using NextDenovo (v2.5.2)^49^. The raw assembly was polished in three rounds with DNBSEQ-T7 short reads via NextPolish (v1.4.0)^50^. Hi-C data were then employed to anchor contigs to chromosomes using HapHiC (v1.0.6)^51^. Manual inspection and adjustment of contig connections and orientations were performed in Juicebox (v1.11.08)^52^, yielding a chromosome-scale genome. Assembly quality was assessed by mapping HiFi reads to the genome with Minimap2^53^ and calculating mapping rate and coverage depth using SAMtools v1.20^54^. Completeness was evaluated with BUSCO v5.7.1 against the embryophyta_odb10 database^55^. Quality value was estimated with Merqury v1.3^56^. MCScan was applied to perform genome synteny analysis^57^.

Prior to gene annotation, repetitive sequences were masked with EDTA v2.0.0^58^. Gene prediction integrated ab initio, homology-based, and transcriptome-supported approaches. Briefly, BRAKER v2.1.6 was employed to train GeneMark-ET and Augustus models using both short-read RNA-seq and PacBio Iso-seq data^59^. Two iterative annotation rounds were conducted with MAKER v3.01.03^60^. In the first round, RNA-seq reads were aligned to the genome using HISAT2 v2.2.1^61^ and assembled with StringTie v2.0^62^, while Iso-seq data were processed via SMRT Analysis Isoseq3. Homology searches were performed using BLAST. The second round incorporated trained gene predictors from SNAP^63^, GeneMark-ET^64^, and Augustus v3.4.0^65^. Final gene models were consolidated with PASA v2.5.1^66^, incorporating alternative splicing variants and untranslated regions. Annotation completeness was assessed using BUSCO v5.7.1.

#### Total RNA isolation, RNA-Seq, and gene expression quantification

Total RNA was extracted with the TransZol Plus RNA Kit (TransGen Biotech, Beijing, China) according to the manufacturer’s instructions. A transcriptome library for RNA-seq was constructed from 1 μg of total RNA using the Illumina Stranded mRNA Prep, Ligation Kit (Illumina, San Diego, USA). Sequencing was conducted on the NovaSeq X Plus platform (Illumina) by Shanghai Majorbio Bio-pharm Biotechnology Co., Ltd. (Shanghai, China).

The raw RNA-seq reads were processed with fastp (v0.23.4)^67^ using default parameters to remove adapter sequences and perform quality control. The resulting high-quality reads were then aligned to the *C. pilosula* reference genome with orientation mode using HISAT2 (v2.2.1)^61^. The mapped reads of each sample were assembled by StringTie (v2.2.1)^62^ in a reference-based approach. The expression level of each transcript was quantified as transcripts per million reads (TPM) using RSEM (v1.3.3)^68^. Differential expression analysis was performed using DESeq2 (v1.42.0)^69^, with genes considered significantly differentially expressed (DEGs) if they met the threshold of |log2FC| ≥ 1 and FDR < 0.05.

#### Comparative genomic analysis

First, the CP2.0_genome sequence was aligned to the reference genome Ref (CP1.0)^25^ using MUMMER (v3.23)^70^ to determine the order and orientation of its contigs. Meanwhile, the potential centromere and telomere regions were systematically predicted using the CentroMiner and TeloExplorer tools within the quarTeT (v1.2.0)^71^ software package under default parameters. Subsequently, the synteny map between the CP2.0 genome and the Ref (CP1.0) genome was generated using the online tool GenomeSyn (v. 1.2)^72^.

#### Phylogenetic analysis

A maximum-likelihood (ML) phylogenetic tree was reconstructed using 72 single-copy orthologous genes identified with OrthoFinder (v2.5.4)^73^ to resolve the phylogenetic relationships among *Vitis vinifera*, *Scutellaria baicalensis*, *Escallonia rubra*, *Lonicera caerulea*, *Aralia elata*, *Daucus carota*, *Centella asiatica*, *Nymphoides indica*, *Scaevola taccada*, *Carthamus tinctorius*, *Lactuca sativa*, *Helianthus annuus*, *Lobelia seguinii*, *P. grandiflorus*, *C. lanceolata*, and *C. pilosula*. The protein sequences were aligned with MAFFT (v7.487)^74^, trimmed using TrimAl (v1.4.rev15)^75^, and subsequently concatenated in PhyloSuite (v1.2.2)^76^. Phylogenetic inference under the maximum-likelihood criterion was performed with RAxML (v8.2.12)^77^, using the PROTGAMMAJTT model for amino acid substitutions and 1000 bootstrap replicates for node support evaluation.

The divergence time of the selected species was estimated using MCMCTREE in PAML (v4.10.7)^78^ under the following settings: 50,000 iterations were run with a sampling frequency of 10, following a burn-in of 200,000 iterations. Fossil-calibrated node constraints were obtained from TimeTree (http://www.timetree.org) and applied as follows: *L. sativa* and *H. annuus* diverged 31.4–38.4 MYA, *C. tinctorius* and *L. sativa* diverged 36.6–45.1 MYA, *S. taccada* and *C. tinctorius* diverged 50.2–79.4 MYA, *S. taccada* and *N. indica* diverged 65.0–90.5 MYA, *C. pilosula* and *C. lanceolata* diverged 6.8–11.4 MYA, *P. grandiflorus* and *C. pilosula* diverged 18.3–37.7 MYA, *E. rubra* and *L. caerulea* diverged 69.0–113 MYA, *D. carota* and *C. asiatica* diverged 18.7–87.4 MYA, *A. elata* and *D. carota* diverged 54.3–69.0 MYA, *S. baicalensis* and *V. vinifera* diverged 111.4–123.9 MYA.

Gene family evolution, including expansion and contraction, was inferred using CAFE (v5.0)^79^. Results were visualized using custom Python scripts and the iTOL online platform (https://itol.embl.de/). Functional enrichment analysis of expanded gene families was conducted based on Gene Ontology (GO) and the Kyoto Encyclopedia of Genes and Genomes (KEGG) databases using ClusterProfiler (v4.0)^80^.

#### Whole-genome duplication analysis

The WGD events analysis method was modified by Xie et al^81^. Protein sequences within a genome or between different genomes were aligned by BLASTP (v2.16.0). Matched genes with e value < 1e−5 were considered as potential homologous genes. Next, syntenic blocks within a genome or between different genomes were determined based on the detected homologous gene pairs using WGDI (v1.1). WGD events were inferred from the syntenic relationships within a genome.

#### Candidates of UDP-glycosyltransferases

To identify UGT genes in *C. pilosula* and the other species included in the phylogenetic analysis, hidden Markov model (HMM) profiles of the UDPGT domain (PF00201) were obtained from Pfam (https://pfam.xfam.org/). Using HMMER v3.1b2^82^, we screened the proteomes of the target species with a significance cutoff of E < 1e-5. Additionally, protein sequences of *A. thaliana* UGTs were retrieved from TAIR (https://www.arabidopsis.org/) and used as queries for BLASTP searches against the protein databases of *C. pilosula* and the comparative species (E < 1e-5). The candidate UGT genes were defined as the intersection of the sequences identified by both methods. Open reading frames (ORFs) were predicted, and sequences shorter than 1000 bp were discarded. For the final screening of UGTs, we identified high-expression DEGs (TPM > 5) in the periderm and generated corresponding heatmaps using the Heatmap Illustrator function in TBtools-II^83^.

#### Identification, phylogenetic, and collinearity analysis of UGT76 members

For the identification of UGT76 family genes in *C. pilosula* and the co-analyzed species, a BLASTP-based comparison was conducted against 157 reference UGT76 sequences from the Plant UDP-glycosyltransferases Database (E-value < 1e-5, amino acid identity > 45%; https://labs.wsu.edu/ugt/). Additionally, the putative UGT protein sequences from *C. pilosula* and the other species were submitted to the Plant Synthetic BioDatabase (https://www.bic.ac.cn/PSBD/front/#/) for functional classification and evolutionary inference of the UGT76 family. The final set of UGT76 genes was determined by integrating the results from both approaches.

To reconstruct the phylogenetic relationships, protein sequences of the identified UGT76 genes from *C. pilosula* and the related species, along with the 157 reference UGT76 sequences, were aligned using MAFFT v7.487^74^ and subsequently refined with TrimAl v1.4^75^. Phylogenetic inference was performed with RAxML-NG v1.0.376^84^ under the best-fit model selected by ModelTest-NG v0.1.7^85^. The UGT85A1 from *A. thaliana* was used as the outgroup.

Furthermore, synteny analysis between each pair of the examined species, including *V. vinifera, S. baicalensis, A. elata, L. seguinii, P. grandiflorus, C. lanceolata, and C. pilosula*, was conducted using MCScan (Python version)^57^. The targeted UGT76 genes or their flanking genes were used as seeds to identify syntenic blocks indicative of evolutionary conservation.

#### Reconstructions of ancestral sequences

Using the aligned sequence file, phylogenetic tree file, and a configured control file, the ancestral sequences of UGT76 family members were inferred under the JTT + GAMMA model using the CodeML program in PAML (v4.10.7)^78^. Regions containing alignment gaps were reconstructed via the parsimony method to determine the most probable ancestral residues. The corresponding ancestral genes were subsequently synthesized after codon optimization and functionally characterized in *E. coli* BL21(DE3). Sequence information of all UGT76 ancestors was shown in Supplementary Data 17.

#### Recombinant protein expression, purification, and enzyme activity assay

Candidate UGT76 genes were amplified with the primers listed in Supplementary Data 20 using KOD DNA polymerase (Toyobo, Osaka, Japan). The resulting PCR products were cloned into the pET-28a (+) vector via the NdeI and NotI restriction sites using the ClonExpress II One Step Cloning Kit (Vazyme, Nanjing, China). The recombinant plasmids were subsequently transformed into *E. coli* BL21 (DE3) (TransGen Biotech, Beijing, China). As a control, the empty pET-28a (+) vector was also transformed into *E. coli* BL21 (DE3). For protein expression, single colonies were inoculated into LB medium supplemented with 50 μg/mL kanamycin and cultured at 37 °C with shaking at 200 rpm. When the OD_600_ reached 0.5–0.6, protein expression was induced by adding 0.1 mM IPTG, followed by incubation at 18 °C for 20 h. Cells were harvested and lysed by sonication on ice. The recombinant proteins were purified using a nickel-affinity column. After SDS-PAGE analysis, the target protein was concentrated and desalted using a 30-kDa ultrafiltration device (Merck Millipore, Billerica, MA, USA) with an appropriate storage buffer.

The reactions were performed in 100 μL mixtures consisting of 50 mM PBS (pH 7.4), 0.1 mM substrate, 1 mM UDP-Glc, and 10 μg of purified protein. After incubation at 37 °C for 2 h, the reactions were stopped by adding an equal volume of ice-cold methanol, followed by centrifugation at 13,000 rpm for 30 min. The resulting supernatants were then subjected to LC-MS analysis.

To evaluate sugar donor specificity, enzymatic reactions were performed as described above using UDP-Galactose, UDP-Glucuronic acid, UDP-Rhamnose, and UDP-Xylose (1 mM each) in place of UDP-Glc. Substrate specificity was further assessed using structurally related compounds: falcarindiol (a C17 natural polyacetylene diol), 10-undecyn-1-ol (a C11 simple alkynol), and 4-pentyn-1-ol (a C5 simple alkynol), each at 0.1 mM final concentration. Reactions were performed under standard conditions and analyzed by LC-MS.

#### Enzyme kinetics of CpUGT76BG1, CpUGT76BG2 and CpUGT94BY2

For steady-state kinetic analysis, reactions were performed in 100 μL mixtures consisting of 50 mM PBS (pH 7.4), varying concentrations of acceptor substrate (0–500 μM), 1 mM UDP-Glc, and purified enzyme at individually optimized amounts: 0.3 μg for CpUGT76BG1, 1 μg for CpUGT76BG2, and 0.1 μg for CpUGT94BY2. Kinetic parameters (*K_m_, k_cat_, k_cat_/K_m_*) were calculated by fitting the initial velocity data to the Michaelis-Menten equation using GraphPad Prism 10.2.0 and shown in Supplementary Fig. 13.

#### Molecular docking and site-directed mutagenesis

Molecular docking and site-directed mutagenesis were performed as described below. The tertiary structures of the candidate UGTs (CpUGT76BG1, CpUGT76BG2, and CpUGT94BY2) were predicted using AlphaFold3 (https://alphafoldserver.com/). Protein models, UDP-Glc, and substrates (lobetyol and lobetyolin) were prepared using AutoDockTools v1.5.6, followed by molecular docking with AutoDock Vina. Structural graphics were rendered with PyMOL 2.4. Site-directed mutants were generated by overlap extension PCR using the primers listed in Supplementary Data 20.

#### Gene functional characterization in *N. benthamiana*

The target genes for expression in *N. benthamiana* were cloned into the pHB vector via seamless assembly using the *Pst* I and *Bam*H I restriction sites. The resulting recombinant plasmids were subsequently introduced into Agrobacterium tumefaciens strain *GV3101*. Single colonies of the transformed *GV3101* were cultured at 28°C with shaking in LB medium supplemented with kanamycin (50 μg/mL), rifampicin (50 μg/mL), and gentamicin (25 μg/mL) until OD_600_ = 0.6. The bacterial cells were collected by centrifugation and resuspended in MMA buffer. The resulting bacterial suspension was infiltrated into leaves of 4–5 week-old *N. benthamiana* plants. For substrate-feeding assays, a solution of 0.2 mM substrate (prepared in MMA buffer) was infiltrated into the leaves on the third day after agroinfiltration. After an additional 4 days of incubation, the infiltrated leaves were harvested, flash-frozen in liquid nitrogen, and extracted for subsequent LC-MS analysis.

#### Plant transformation and qRT-PCR-analysis

The *CpUGT76BG1*, *CpUGT76BG2*, and *CpUGT94BY2*, respectively, overexpression binary vector was derived from a modified pCAMBIA1305, with the incorporation RUBY as the reporter, driven by the CaMV 35S promoter. All of the plasmids were introduced into *A. rhizogenes* K599 cells through the heat-shock transformation method according to the manufacturer (AC1080; WEIDI, China). Four weeks old seedlings of *C. pilosula* were selected for transformation and all the steps following were performed with the previously described method by Cao et al^86^. All of the PCR primers were listed in Supplementary Data 20. After collecting red positive hairy roots from overexpression lines and colorless hairy roots from wild-type lines individually per plant, the expression level of the target gene and the contents of lobetyolin and lobetyolinin were analyzed separately in the overexpression and wild-type lines. The quantification of lobetyolin and lobetyolinin was performed through calibration curves with authentic standards (Supplementary Fig. 14). Results were normalized to sample dry weight (mg. g-1 DW) of *C. pilosula* (Supplementary Data 14). Data were acquired by the software Masslynx 4.2.

## Data availability

The whole genome sequence data reported in this paper have been deposited in the Genome Warehouse in National Genomics Data Center, Beijing Institute of Genomics, Chinese Academy of Sciences/China National Center for Bioinformation, under accession number PRJCA047761 that is publicly accessible at https://ngdc.cncb.ac.cn/gwh. The transcriptome data of *C. pilosula* from different root tissues reported in this paper have been deposited in the Genome Sequence Archive in National Genomics Data Center, Beijing Institute of Genomics, Chinese Academy of Sciences (GSA: CRA031669) that are publicly accessible at https://ngdc.cncb.ac.cn/gsa.

## Acknowledgements

The author the UGT Nomenclature Committee to acquire its specific designation for naming of the *CpUGT76BG1, CpUGT76BG2 and CpUGT94BY2* (https://labs.wsu.edu/ugt/).

## Author contributions

Conceptualization, S.Q., W.S.C., J.D.H., and W.S.; Methodology, J.D.H., R.J.M., X.C., and Z.C.X.; Validation, S.Q., J.D.H., X.C., and C.H.W.; Formal Analysis, J.D.H., C.H.W., S.S.C., S.Q., and C.Z.; Investigation, S.Q., J.D.H., X.C., S.S.C., and W.S.C.; Writing – Original Draft, S.Q., J.D.H., W.S., and X.C.; Writing – Review & Editing, S.Q., J.D.H., W.S.C., and X.C.; Resources, W.S.C., P.D., J.D.H., Y.X., and R.J.M.; Supervision, W.S.C.; Funding Acquisition, W.S.C., S.Q., Y.X., W.S., J.D.H., and R.J.M. The authors J.D.H., M.H., X.C., and C.H.W., contributed equally to this work.

## Additional information

## References

1. Wang, S., Alseekh, S., Fernie, A.R. & Luo, J. The Structure and Function of Major Plant Metabolite Modifications. Mol. Plant 12, 899–919 (2019).

2. Lemke, O. et al. The role of metabolism in shaping enzyme structures over 400 million years. Nature 644, 280–289 (2025).

3. Lee, Y.J., Cho, Y. & Tran, H.N.K. Secondary Metabolites from the Marine Sponges of the Genus *Petrosia*: A Literature Review of 43 Years of Research. Mar. Drugs 19, 122 (2021).

4. Li, J., et al. Research progress on chemistry, bioactivity, and NMR patterns of acetylene compounds in natural products (2015–2025). Phytochem. Rev. 25, 3281–3319 (2026).

5. Lai, J.X. et al. Plant polyacetylenoids: Phytochemical, analytical and pharmacological updates. *Arab*. J. Chem. 16, 105137 (2023).

6. Pollo, L.A.E., Frederico, M.J., Bortoluzzi, A.J., Silva, F. & Biavatti, M.W. A new polyacetylene glucoside from *Vernonia scorpioides* and its potential antihyperglycemic effect. Chem. Biol. Interact. 279, 95–101 (2018).

7. Santos, J.A.M., Santos, C., Freitas Filho, J.R., Menezes, P.H. & Freitas, J.C.R. Polyacetylene Glycosides: Isolation, Biological Activities and Synthesis. Chem. Rec. 22, e202100176 (2022).

8. Xie, Q. & Wang, C. Polyacetylenes in herbal medicine: A comprehensive review of its occurrence, pharmacology, toxicology, and pharmacokinetics (2014-2021). Phytochemistry 201, 113288 (2022).

9. Zhang, S., Chai, X., Hou, G., Zhao, F. & Meng, Q. Platycodon grandiflorum (Jacq.) A. DC.: A review of phytochemistry, pharmacology, toxicology and traditional use. Phytomedicine 106, 154422 (2022).

10. Yue, J., Xiao, Y. & Chen, W. Insights into Genus *Codonopsis*: From past Achievements to Future Perspectives. Crit. Rev. Anal. Chem. 54, 3345–3376 (2024).

11. Folquitto, D.G. et al. Biological activity, phytochemistry and traditional uses of genus *Lobelia* (Campanulaceae): A systematic review. Fitoterapia 134, 23–38 (2019).

12. Santos, J.A.M. et al. Synthesis, and antitumoral and antiviral evaluation of polyacetylene glycoside derivatives. Org. Biomol. Chem. 23, 410–421 (2025).

13. Zhou, J. et al. An inulin-type fructan CP-A from Codonopsis pilosula attenuates experimental colitis in mice by promoting autophagy-mediated inactivation of NLRP3 inflammasome. Chin. J. Nat. Med. 22, 249–264 (2024).

14. Yonekura-Sakakibara, K. & Hanada, K. An evolutionary view of functional diversity in family 1 glycosyltransferases. Plant J. 66, 182–193 (2011).

15. De Bruyn, F., Maertens, J., Beauprez, J., Soetaert, W. & De Mey, M. Biotechnological advances in UDP-sugar based glycosylation of small molecules. Biotechnol. Adv. 33, 288–302 (2015).

16. Yang, T. et al. Hydrophobic recognition allows the glycosyltransferase UGT76G1 to catalyze its substrate in two orientations. Nat. Commun. 10, 3214 (2019).

17. Liu, S.J. et al. Deciphering the biosynthetic pathway of triterpene saponins in *Prunella vulgaris*. Plant J. 121, e17220 (2025).

18. Wang, J. et al. Evolution and functional divergence of glycosyltransferase genes shaped the quality and cold tolerance of tea plants. Plant Cell 37, koae268 (2024).

19. Li, P. et al. Multiomics analyses of two *Leonurus* species illuminate leonurine biosynthesis and its evolution. Mol. Plant 17, 158–177 (2024).

20. Zou, J.L. et al. Complete biosynthetic pathway of furochromones in *Saposhnikovia divaricata* and its evolutionary mechanism in *Apiaceae* plants. Nat. Commun. 16, 3109 (2025).

21. Kim, C.Y. et al. Tracing the stepwise Darwinian evolution of a plant halogenase. Sci. Adv. 11, eadv6898 (2025).

22. An, Z. et al. Lineage-Specific CYP80 Expansion and Benzylisoquinoline Alkaloid Diversity in Early-Diverging Eudicots. Adv. Sci. (Weinh) 11, e2309990 (2024).

23. Tang, W., Shi, J.J., Liu, W., Lu, X. & Li, B. MALDI Imaging Assisted Discovery of a Di-O-glycosyltransferase from Platycodon grandiflorum Root. Angew. Chem. Int. Ed. Engl. 62, e202301309 (2023).

24. Wang, Z.L. et al. Elucidation of the biosynthetic pathway of hydroxysafflor yellow A. Nat. Commun. 16, 4489 (2025).

25. Chen, B.Z. et al. Chromosome-scale genome assembly of *Codonopsis pilosula* and comparative genomic analyses shed light on its genome evolution. Front. Plant. Sci. 15, 1469375 (2024).

26. Liu, Y. et al. pUGTdb: A comprehensive database of plant UDP-dependent glycosyltransferases. Mol. Plant 16, 643–646 (2023).

27. Osmani, S.A., Bak, S., Imberty, A., Olsen, C.E. & Møller, B.L. Catalytic Key Amino Acids and UDP-Sugar Donor Specificity of a Plant Glucuronosyltransferase, UGT94B1: Molecular Modeling Substantiated by Site-Specific Mutagenesis and Biochemical Analyses. Plant Physiol. 148, 1295–1308 (2008).

28. Jiang, Z. et al. Molecular characterization and structural basis of a promiscuous glycosyltransferase for beta-(1,6) oligoglucoside chain glycosides biosynthesis. Plant Biotechnol. J. 23, 2242–2253 (2025).

29. Sun, W. et al. Pharmacokinetic profiles of falcarindiol and oplopandiol in rats after oral administration of polyynes extract of *Oplopanax elatus*. Chin. J. Nat. Med. 14, 714–720 (2016).

30. Li, C. et al. Single-cell multi-omics in the medicinal plant *Catharanthus roseus*. Nat. Chem. Biol. 19, 1031–1041 (2023).

31. Busta, L. et al. Identification of Genes Encoding Enzymes Catalyzing the Early Steps of Carrot Polyacetylene Biosynthesis. Plant Physiol. 178, 1507–1521 (2018).

32. Zhu, J.J. et al. Divergent fatty acid desaturase 2 is essential for falcarindiol biosynthesis in carrot. Plant Commun. 6, 101323 (2025).

33. Jeon, J.E. et al. A Pathogen-Responsive Gene Cluster for Highly Modified Fatty Acids in Tomato. Cell 180, 176–187 (2020).

34. Scott, S., Cahoon, E.B. & Busta, L. Variation on a theme: the structures and biosynthesis of specialized fatty acid natural products in plants. Plant J. 111, 954–965 (2022).

35. Minto, R.E. & Blacklock, B.J. Biosynthesis and function of polyacetylenes and allied natural products. Prog. Lipid Res. 47, 233–306 (2008).

36. Santos, P., Busta, L., Yim, W.C., Cahoon, E.B. & Kosma, D.K. Structural diversity, biosynthesis, and function of plant falcarin-type polyacetylenic lipids. J. Exp. Bot. 73, 2889–2904 (2022).

37. Wang, H.T. et al. Insights into the Mechanisms of Sugar Acceptor Selectivity of Plant Flavonoid Apiosyltransferases. J. Am. Chem. Soc. 147, 20631–20643 (2025).

38. Chen, Y. et al. Unraveling the serial glycosylation in the biosynthesis of steroidal saponins in the medicinal plant *Paris polyphylla* and their antifungal action. Acta Pharm. Sin. B 13, 4638–4654 (2023).

39. Liu, Z.F., Li, J.X., Sun, Y.W., Zhang, P. & Wang, Y. Structural Insights into the Catalytic Mechanism of a Plant Diterpene Glycosyltransferase SrUGT76G1. Plant Commun. 1, 100004 (2020).

40. Li, Y., Winzer, T., He, Z. & Graham, I.A. Over 100 Million Years of Enzyme Evolution Underpinning the Production of Morphine in the Papaveraceae Family of Flowering Plants. Plant Commun. 1, 100029 (2020).

41. Qiu, S. et al. Functional evolution and diversification of CYP82D subfamily members have shaped flavonoid diversification in the genus *Scutellaria*. Plant Commun. 6, 101134 (2025).

42. Ishimaru, K., Yonemitsu, H. & Shimomura, K. Lobetyolin and lobetyol from hairy root culture of *Lobelia inflata*. Phytochemistry 30, 2255–2257 (1991).

43. Ishimaru, K. et al. Polyacetylene glycosides from *Pratia nummularia* cultures. Phytochemistry 62, 643–646 (2003).

44. Lieberman-Aiden, E. et al. Comprehensive mapping of long-range interactions reveals folding principles of the human genome. Science 326, 289–293 (2009).

45. Marcais, G. & Kingsford, C. A fast, lock-free approach for efficient parallel counting of occurrences of k-mers. Bioinformatics 27, 764–770 (2011).

46. Vurture, G.W. et al. GenomeScope: fast reference-free genome profiling from short reads. Bioinformatics 33, 2202–2204 (2017).

47. Cheng, H., Concepcion, G.T., Feng, X., Zhang, H. & Li, H. Haplotype-resolved de novo assembly using phased assembly graphs with hifiasm. Nat. Methods 18, 170–175 (2021).

48. Guan, D. et al. Identifying and removing haplotypic duplication in primary genome assemblies. Bioinformatics 36, 2896–2898 (2020).

49. Hu, J. et al. NextDenovo: an efficient error correction and accurate assembly tool for noisy long reads. Genome Biol. 25, 107 (2024).

50. Hu, J., Fan, J., Sun, Z. & Liu, S. NextPolish: a fast and efficient genome polishing tool for long-read assembly. Bioinformatics 36, 2253–2255 (2020).

51. Zeng, X. et al. Chromosome-level scaffolding of haplotype-resolved assemblies using Hi-C data without reference genomes. Nat. Plants 10, 1184–1200 (2024).

52. Durand, N.C. et al. Juicebox Provides a Visualization System for Hi-C Contact Maps with Unlimited Zoom. Cell Syst. 3, 99–101 (2016).

53. Li, H. Minimap2: pairwise alignment for nucleotide sequences. Bioinformatics 34, 3094–3100 (2018).

54. Li, H. et al. The Sequence Alignment/Map format and SAMtools. Bioinformatics 25, 2078–2079 (2009).

55. Manni, M., Berkeley, M.R., Seppey, M., Simao, F.A. & Zdobnov, E.M. BUSCO Update: Novel and Streamlined Workflows along with Broader and Deeper Phylogenetic Coverage for Scoring of Eukaryotic, Prokaryotic, and Viral Genomes. Mol. Biol. Evol. 38, 4647–4654 (2021).

56. Rhie, A., Walenz, B.P., Koren, S. & Phillippy, A.M. Merqury: reference-free quality, completeness, and phasing assessment for genome assemblies. Genome Biol. 21, 245 (2020).

57. Tang, H., et al. JCVI: A versatile toolkit for comparative genomics analysis. Imeta 3, e211 (2024).

58. Ou, S. et al. Benchmarking transposable element annotation methods for creation of a streamlined, comprehensive pipeline. Genome Biol. 20, 275 (2019).

59. Bruna, T., Hoff, K.J., Lomsadze, A., Stanke, M. & Borodovsky, M. BRAKER2: automatic eukaryotic genome annotation with GeneMark-EP+ and AUGUSTUS supported by a protein database. NAR Genom. Bioinform. 3, lqaa108 (2021).

60. Cantarel, B.L. et al. MAKER: an easy-to-use annotation pipeline designed for emerging model organism genomes. Genome Res. 18, 188–196 (2008).

61. Kim, D., Langmead, B. & Salzberg, S.L. HISAT: a fast spliced aligner with low memory requirements. Nat. Methods 12, 357–360 (2015).

62. Pertea, M. et al. StringTie enables improved reconstruction of a transcriptome from RNA-seq reads. Nat. Biotechnol. 33, 290–295 (2015).

63. Korf, I. Gene finding in novel genomes. BMC Bioinformatics 5, 59 (2004).

64. Lomsadze, A., Ter-Hovhannisyan, V., Chernoff, Y.O. & Borodovsky, M. Gene identification in novel eukaryotic genomes by self-training algorithm. Nucleic Acids Res. 33, 6494–6506 (2005).

65. Stanke, M., Tzvetkova, A. & Morgenstern, B. AUGUSTUS at EGASP: using EST, protein and genomic alignments for improved gene prediction in the human genome. Genome Biol. 7 **Suppl 1**, S11.1-S11.8 (2006).

66. Haas, B.J. et al. Improving the *Arabidopsis* genome annotation using maximal transcript alignment assemblies. Nucleic Acids Res. 31, 5654–5666 (2003).

67. Chen, S., Zhou, Y., Chen, Y. & Gu, J. fastp: an ultra-fast all-in-one FASTQ preprocessor. Bioinformatics 34, i884–i890 (2018).

68. Li, B. & Dewey, C.N. RSEM: accurate transcript quantification from RNA-Seq data with or without a reference genome. BMC Bioinformatics 12, 323 (2011).

69. Love, M.I., Huber, W. & Anders, S. Moderated estimation of fold change and dispersion for RNA-seq data with DESeq2. Genome Biol. 15 550 (2014).

70. Kurtz, S. et al. Versatile and open software for comparing large genomes. Genome Biol. 5, R12 (2004).

71. Lin, Y. et al. quarTeT: a telomere-to-telomere toolkit for gap-free genome assembly and centromeric repeat identification. Hortic. Res. 10, uhad127 (2023).

72. Zhou, Z.W. et al. GenomeSyn: a bioinformatics tool for visualizing genome synteny and structural variations. J. Genet. Genomics 49, 1174–1176 (2022).

73. Emms, D.M. & Kelly, S. OrthoFinder: phylogenetic orthology inference for comparative genomics. Genome Biol. 20 238 (2019).

74. Katoh, K. & Standley, D.M. MAFFT Multiple Sequence Alignment Software Version 7: Improvements in Performance and Usability. Mol. Biol. Evol. 30, 772–780 (2013).

75. Capella-Gutierrez, S., Silla-Martinez, J.M. & Gabaldon, T. trimAl: a tool for automated alignment trimming in large-scale phylogenetic analyses. Bioinformatics 25, 1972–1973 (2009).

76. Zhang, D. et al. PhyloSuite: An integrated and scalable desktop platform for streamlined molecular sequence data management and evolutionary phylogenetics studies. Mol. Ecol. Resour. 20, 348–355 (2020).

77. Stamatakis, A. RAxML version 8: a tool for phylogenetic analysis and post-analysis of large phylogenies. Bioinformatics 30, 1312–1313 (2014).

78. Yang, Z.H. PAML 4: Phylogenetic analysis by maximum likelihood. Mol. Biol. Evol. 24, 1586–1591 (2007).

79. Mendes, F.K., Vanderpool, D., Fulton, B. & Hahn, M.W. CAFE 5 models variation in evolutionary rates among gene families. Bioinformatics 36, 5516–5518 (2021).

80. Wu, T. et al. clusterProfiler 4.0: A universal enrichment tool for interpreting omics data. Innovation 2, 100141 (2021).

81. Xie, D. et al. The wax gourd genomes offer insights into the genetic diversity and ancestral cucurbit karyotype. Nat. Commun. 10, 5158 (2019).

82. Mistry, J., Finn, R.D., Eddy, S.R., Bateman, A. & Punta, M. Challenges in homology search: HMMER3 and convergent evolution of coiled-coil regions. Nucleic Acids Res. 41, e121 (2013).

83. Chen, C. et al. TBtools-II: A “one for all, all for one” bioinformatics platform for biological big-data mining. Mol. Plant 16, 1733–1742 (2023).

84. Kozlov, A.M., Darriba, D., Flouri, T., Morel, B. & Stamatakis, A. RAxML-NG: a fast, scalable and user-friendly tool for maximum likelihood phylogenetic inference. Bioinformatics 35, 4453–4455 (2019).

85. Darriba, D. et al. ModelTest-NG: A New and Scalable Tool for the Selection of DNA and Protein Evolutionary Models. Mol. Biol. Evol. 37, 291–294 (2020).

86. Cao, X. et al. A simple widely applicable hairy root transformation method for gene function studies in medicinal plants. Acta Pharm. Sin. B 15, 4300–4305 (2025).

